# Structure-Kinetics Relationships of Opioids from Metadynamics and Machine Learning

**DOI:** 10.1101/2023.03.06.531338

**Authors:** Paween Mahinthichaichan, Ruibin Liu, Quynh N. Vo, Christopher R. Ellis, Lidiya Stavitskaya, Jana Shen

**Affiliations:** Center for Drug Evaluation and Research, United States Food and Drug Administration, Silver Spring, MD 20993, United States; Department of Pharmaceutical Sciences, University of Maryland School of Pharmacy, Baltimore, MD 21201, United States; ComputChem LLC, Baltimore, MD 21230; United States Army, DEVCOM Chemical Biological Center, Aberdeen Proving Ground, MD 21010, United States

## Abstract

The nation’s opioid overdose deaths reached an all-time high in 2021. The majority of deaths are due to synthetic opioids represented by fentanyl. Naloxone, which is a FDA-approved reversal agent, antagonizes opioids through competitive binding at the *μ*-opioid receptor (mOR). Thus, knowledge of opioid’s residence time is important for assessing the effectiveness of naloxone. Here we estimated the residence times of 15 fentanyl and 4 morphine analogs using metadynamics, and compared them with the most recent measurement of the opioid kinetic, dissociation, and naloxone inhibitory constants (Mann, Li et al, Clin. Pharmacol. Therapeut. 2022). Importantly, the microscopic simulations offered a glimpse at the common binding mechanism and molecular determinants of dissociation kinetics for fentanyl analogs. The insights inspired us to develop a machine learning (ML) approach to analyze the kinetic impact of fentanyl’s substituents based on the interactions with mOR residues. This proof-of-concept approach is general; for example, it may be used to tune ligand residence times in computeraided drug discovery.

**Graphical TOC Entry:** 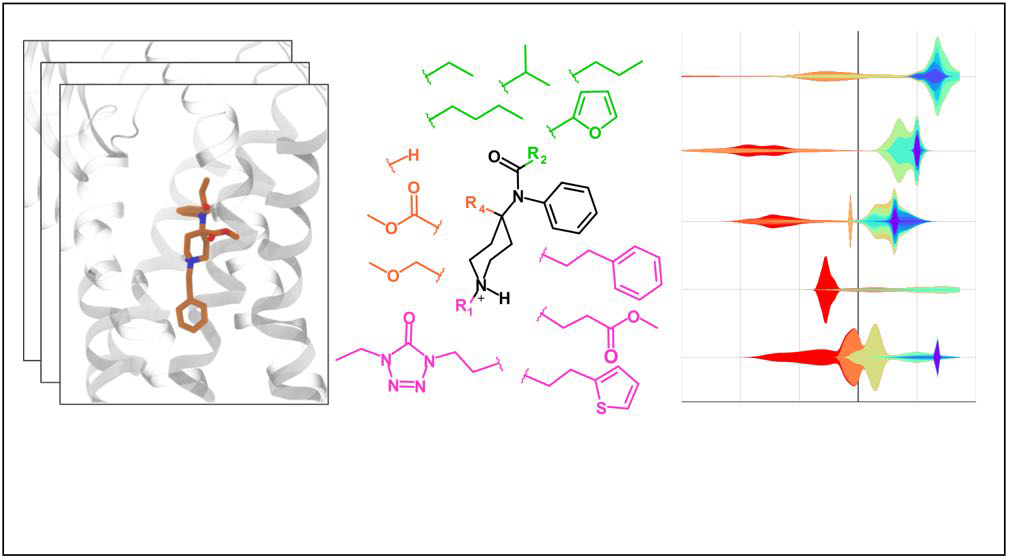

## INTRODUCTION

In 2021, the nation set a new record of over 100,000 deaths from illicit and prescription drug overdose, (https://www.cdc.gov/nchs/pressroom/nchs_press_releases/2021/20211117.htm). The primary cause of overdose-related death is respiratory depression.^1^ At a molecular level, fentanyl and other opioids are agonist of the *μ*–opioid receptor (mOR, Fig. 1a), which is a member of the class A family of G-protein coupled receptors (GPCRs).^2^ Upon fentanyl binding, mOR undergoes a conformational change, allowing it to associate with G-proteins and *β*-arrestins, which in turn activates the related signaling pathways. It is hypothesized that the G-protein pathway produces analgesic effect while the *β*-arrestin pathway is mainly responsible for adverse effects such as respiratory depression and constipation.^3,4^ Compared to morphine (a naturally occurring opiate), fentanyl (a synthetic opioid) is 50–400 times more potent and has a faster onset and shorter duration of action.^3,4^ Distinctive from morphine, fentanyl’s core structure is *N*-phenylpiperidin-4-amine (also known as 4-anilidopiperidine), which has 5 positions that can be readily modified to produce various analogs (Fig. 1b and Fig. 1d) with significantly enhanced potencies.^5,6^ For example, a methyl ester substitution at the R4 position produces carfentanil (Fig. 1d), which is 20–100 times more potent than fentanyl.^3,7^ Carfentanil, which is approved for large animal veterinary medicine, ^3,7^ is the most potent opioid used commercially. Although not approved for humans, carfentanil is a known additive to other drugs of abuse in the United States and Canada, contributing to overdose-related deaths. ^7^ Starting from carfentanil, a methyl substitution at the R5 produces even more potent and long-lasting lofentanil (Fig. 1d), which is placed in the schedule I controlled substance list by the US Drug Enforcement Agency (https://www.deadiversion.usdoj.gov/schedules/#abc) with no currently accepted medical use and a high potential for abuse. Due to the ease and low cost of chemical synthesis, a large number of fentanyl analogs have been made and sold on the darknet market in various forms, exacerbating the opioid crisis.^8,9^

**Figure 1.**
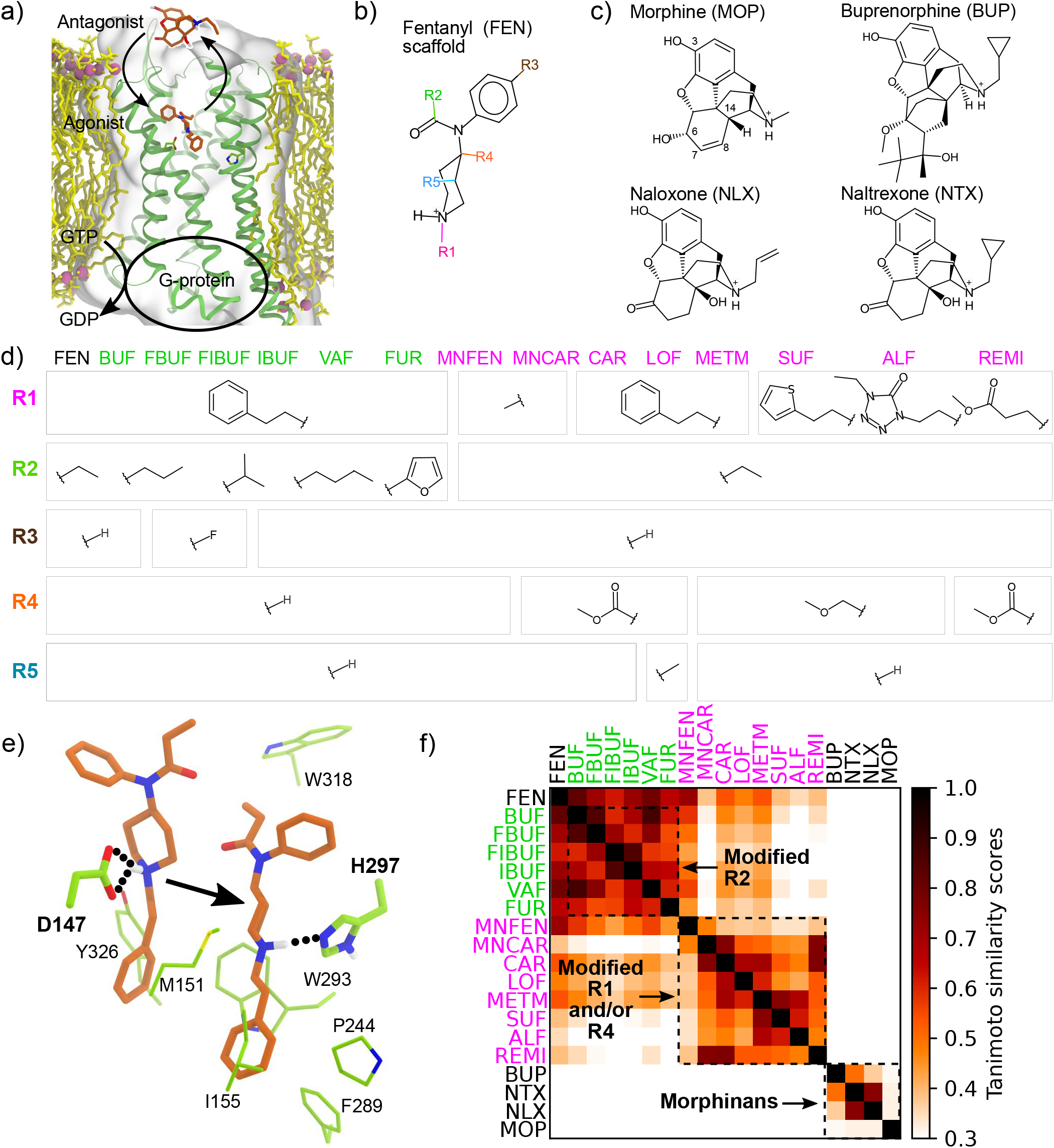
The fentanyl and morphine analogs studied in this work. **a)** Schematic view of the structure and function of *μ*-opioid receptor (mOR). **b)** The fentanyl (FEN) scaffold with the commonly modified substituent groups. **c)** The structures of morphine (MOP) and analogs. **d)** The substituent groups for the 15 fentanyl analogs studied here. The names of compounds that mainly differ in R2 are in green, and those that mainly differ in R1 and/or R4 are in magenta. The abbreviations are explained in the main text. **e)** The two putative binding modes of fentanyl analogs. **f)** Tanimoto similarity matrix of the 19 fentanyl and morphine analogs.

Currently, naloxone is the most widely used rescue agent for opioid overdose approved by the U.S. Food and Drug Administration (FDA).^9^ Although naloxone is generally considered effective, recent data suggest that higher or multiple dosing strategies are needed in the era of continued abuse of fentanyl and the availability of analogs with even higher potencies.^6,9,10^ At a molecular level, naloxone reverses opioid-induced respiratory depression through competitive binding and antagonizing of mOR (Fig. 1a). Therefore, it is plausible that opioids with slower dissociation kinetics may be difficult to reverse. Indeed, a most recent study of opioid-receptor binding and dissociation indicated that carfentanil, which was difficult to reverse in the virtual patient simulation, has a substantially slower dissociation rate constant as compared to the tested fentanyl and derivatives.^11^ Thus, obtaining kinetics data may be of significant value to the evaluation of naloxone’s effectiveness for preventing overdose related deaths. However, measurement of dissociation rate is very costly and time consuming due to the use of radioligand labeling. Considering the large number of existing fentanyl analogs (more than 1400 according to a recent article ^8^) and continually emerging new ones on the darknet market, there is an urgent need to develop a predictive capability that can evaluate the dissociation kinetics of a large number of opioids.

Recognizing the aforementioned need, we recently implemented a computational protocol^12^ based on a biased sampling molecular dynamics (MD) method called infrequent well-tempered metadynamics ^13–15^ and X-ray structure of the agonist BU72-bound mOR^16^ to estimate fentanyl’s residence time (inverse of dissociation rate constant *k_off_*). This work^12^ and another study^17^ based on the weighted-ensemble (WE) MD (an unbiased enhanced sampling MD protocol)^18^ demonstrated that, in addition to the canonical binding mode, in which fentanyl’s piperidine amine forms a salt bridge with the absolutely conserved D147^3.32^ (superscript refers to the Ballesteros-Weinstein numbering^19^) as in the BU72-bound mOR structure, fentanyl can adopt an alternative binding mode, whereby the piperidine’s amine donates a hydrogen bond to the conserved H297^6.52^ when it is in the neutral N*δ*-protonated state (Hid). Related to the binding mode dependence on the H297^6.52^ protonation state, the calculated residence time (*τ*_calc_) of fentanyl with Hid (38±19 s) is more than one order of magnitude larger than with Hie (N_*ϵ*_-protonated) or Hip (doubly protonated) form.^12^ For the sake of clarity, we will drop the Ballesteros-Weinstein numbering in the discussion that follows. The calculated *τ* with Hid297 is about one order of magnitude smaller than the recent experimental value of 4 mins^11^), which is not surprising, given that the simulation time scale is about 8 orders of magnitude smaller and neither the transition or unbound state regions are thoroughly sampled.^20^

Although metadynamics protocols have been applied to estimate ligand-protein dissociation kinetics in a large number of studies (e.g., in Refs^12,15,20,21^), the prediction accuracy for predicting the relative order of residence times of a large number of structurally related ligands is unclear;^22^ this task is practically more relevant than predicting the absolute residence time of a single ligand. Here we tested the aforementioned metadynamics protocol ^12^ to estimate the residence times of 15 fentanyl and 4 morphine analogs (Fig. 1b–d) and correlated the calculations with the newly obtained experimental estimates for 12 compounds (10 fentanyls and 2 morphinans).^11^ The Pearson’s *r* value between calculation and experiment is 0.65 and it increases to 0.82 when the highly similar R2 modified compounds (mainly derived from butyrfentanyl) are excluded.^11^ Interestingly, the correlation between the calculated *τ*’s and the measured dissociation constants as well as the naloxone inhibitory constants is strong, with the Pearson’s *r* values of −0.87 and 0.90, respectively. Importantly, the metadynamics trajectories allowed us to explore the common binding mechanism and structural modulators of dissociation kinetics. Finally, we developed a machine learning analysis as a proof-of-concept for elucidating the structure-kinetics relationships of opioids based on MD data.

## RESULTS AND DISCUSSION

### Structural analysis of the fentanyl and morphine analogs

A total of 19 fentanyl and morphine analogs are studied in this work (Fig. 1b-d). The fentanyls include butyrfentanyl (BUF), 4-flurobutyrlfentanyl (FBUF), 4-fluoroisobutyrfentanyl (FIBUF), isobutyrfentanyl (IBUF), valerylfentanyl (VAF), furanylfentanyl (FUR), *N*-methyl norfentanyl (MNFEN), N-methyl norcarfentanil (MNCAR), carfentanil (CAR), lofentanil (LOF), 4-methoxymethylfentanyl (METM), sufentanil (SUF), alfentanil (ALF), and remifentanil (REMI). While SUF, REMI and ALF are used in postoperative pain management,^3,6^ LOF, BUF, IBUF, VAF, FUR, FBUF and FIBUF are schedule I controlled substances with no approved medical use (https://www.ecfr.gov/current/title-21/chapter-II/part-1308). Based on the substitutions at the four commonly modified sites on the fentanyl core (Fig. 1b), the fentanyl analogs can be divided into several groups. Compounds in the first group differ from fentanyl mainly in the R2 substitution and they include BUF, FBUF, FIBUF, IBUF, VAF, and FUR (Fig. 1b and d). Due to the small variation of R2, these compounds are similar to fentanyl and to each other, with the Tanimoto similarity scores of between 0.55 and 0.87 (Fig. 1f). Compounds in the second group differ from fentanyl in the R1 and/or R4 substitution, and they include MNFEN, MNCAR, CAR, LOF, METM, SUF, ALF, and REMI. MNFEN is highly similar to fentanyl; the only difference is the phenethyl to methyl substitution at R1 (Fig. 1d and f). CAR, LOF, and METM have a methyl ester or methyl ether substitution at R4 (Fig. 1d). MCF, SUF, ALF, and REMI have both R4 and R1 substitutions (Fig. 1d). As evident from the low (below 0.3) Tanimoto scores, the compounds with R4 or both R4 and R1 substitutions have low structural similarity to fentanyl, although they can be very similar among themselves (Fig. 1f).

The morphinans studied here include morphine (MOP) and the 7,8-dihydromorphine analogs buprenorphine (BUP), naloxone (NLX), and naltrexone (NTX, Fig. 1c). While MOP is an agonist and BUP is a partial agonist, NLX and NTX are antagonists. BUP has modifications at position 6 and 7 as well as a cyclopropylmethyl substitution for the N-methyl group. NLX and NTX have a keto group at position 6 and a hydroxyl substitution at position 14. Additionally, they have allyl or cyclopropylmethyl substitution for the N-methyl group, respectively.

### Estimation of the residence times of fentanyls and morphinans

For each of the 19 fentanyl and morphine analogs, 15 well-tempered infrequent metadynamics simulations ^13,14,23^ were conducted, whereby the starting structures were those relaxed from the modified snapshot of the global free energy minimum from the FEN-mOR WE simulations ^17^ or the docked structure with the X-ray structures of mOR in complex with the agonist BU72 (PDB 5C1M)^16^ or antagonist *β*-FNA (PDB 4DKL, see Methods).^24^ Following our previous work,^12^ a time-dependent biasing potential was deposited every 10 ps (adopted from the work of Casasnovas et al.^20^) along two collective variables, the ligand z position relative to the center of mass of the orthosteric binding site and the number of ligand-mOR contacts. Note, longer deposition times (20 and 30 ps) were tested in our previous work and did not lead to significant different residence time for fentanyl.^12^ The unbinding events (defined as z >15 Å, see justification in our previous work^12^) were observed within 15–90 ns. A total of 285 trajectories were collected, with an aggregate simulation time of ~12 *μ*s. In all simulations, H297 was set to the N_*δ*_ protonated tautomer (Hid) form, as it gave a much slower kinetics and better agreement with experiment for fentanyl than the alternative Hie state.^12^ Following the protocol of Parrinello and coworkers,^20,25^ we fit the empirical cumulative distribution function (CDF) of the rescaled (unbiased) dissociation times recovered from metadynamics trajectories to the theoretical CDF of a homogeneous Poisson process to obtain the characteristic time *τ*_cal_. The two-sample Kolmogorov-Smirnov (KS) test was then used to test the null hypothesis that the sample of dissociation times extracted from metadynamics and a large sample of times randomly generated according to the theoretical CDF reflect the same underlying Poisson distribution. ^25^ The null hypothesis is accepted if the *p*-value is greater than a threshold, which is typically 0.05. A complete description of the methods and protocols is given in SI.

### Correlation between the calculated residence times and related experiment data

Concurrent with the computational study, our colleagues at FDA and the Portland Veterans Affairs Medical Center have measured the association and dissociation rate constants of 12 (out of the aforementioned 19) compounds using radio-labeled ligand-binding assays.^11^ These compounds include BUP, NLX, and 10 fentanyl derivatives: FEN, the R2 modified BUF, FBUF, IBUF, FUR, FIBUF, and the R4 and/or R1 modified CAR, SUF, ALF, and REMI. From the kinetic rate constants, the residence times *τ*_exp_ (1/*k*_off_) and dissociation constants *K_d_* (*k*_off_/*k*_on_) were estimated (Table S1). Additionally, the naloxone inhibitory constant (*K*_i,NLX_) (naloxone concentration that produces 50% reduction in the maximal agonist occupancy in mOR) for a subset (10) of the 12 compounds were also obtained^11^ (Table S1).

A comparison of the calculated and experimental *τ* values (on the log scale) gave a Pearson’s correlation coefficient (*r*) of 0.61 and revealed an overall underestimation of *τ*’s (Fig. 2a and b, Table S1). The latter is not surprising, given the short simulation time and perhaps the different definitions for dissociation. We should also point out that some of experimental estimates, i.e., for FIBUF and FBUF have larger errors.^11^ FIBUF and FBUF belong to the R2 modified compounds which are very similar to fentanyl; thus, it is likely that neither experiment nor simulation is able to accurately quantify the differences in *τ* values for these compounds. Indeed, no correlation was found between the calculated and experimental *τ*’s (Fig. S2). Separating out the R2 modified compounds, the correlation between the experimental and calculated *τ*’s is improved, with the Pearson’s *r* increased to 0.83 (Fig. 2a).

**Figure 2.**
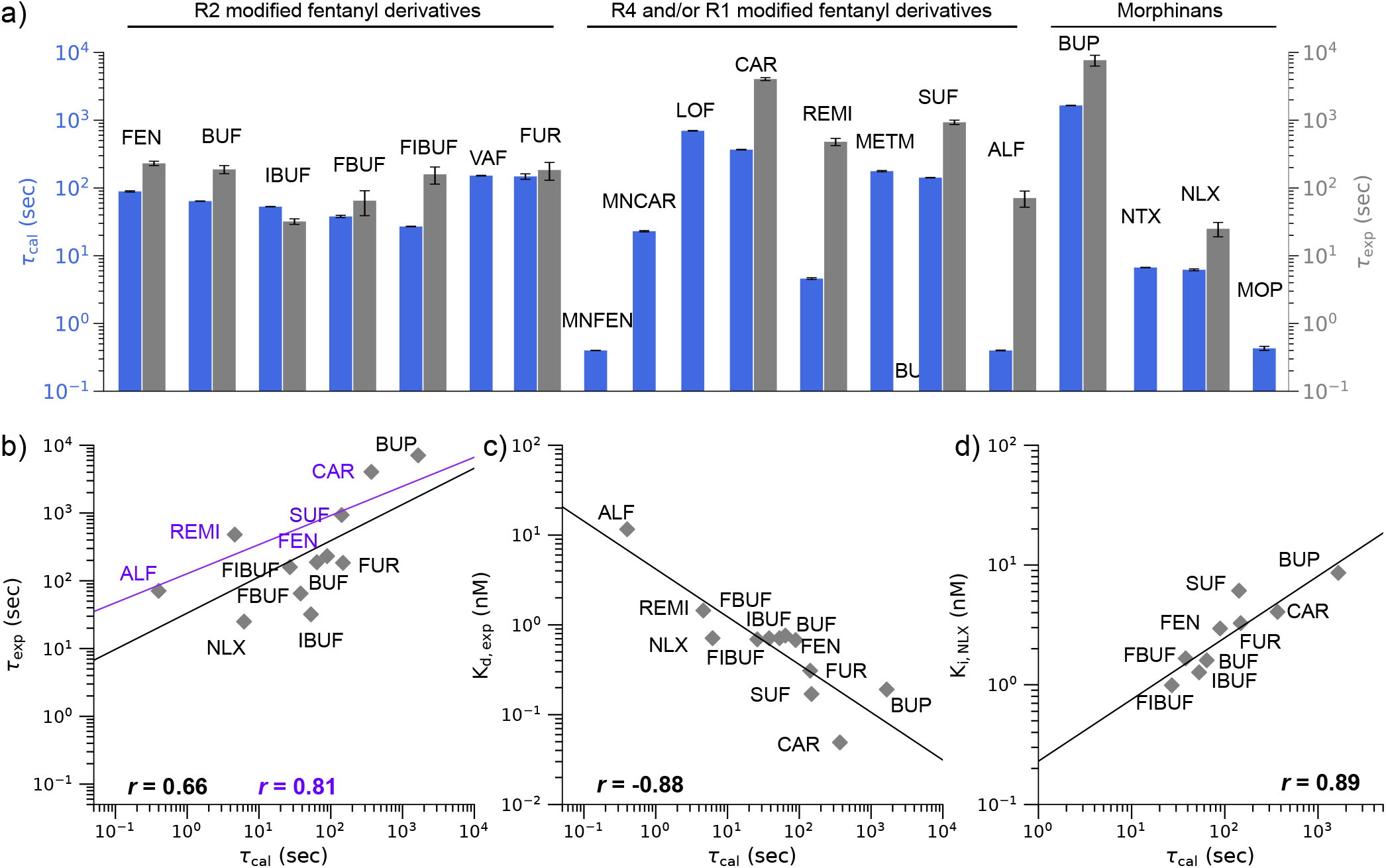
Comparison between the calculated residence times of opioids and related experimental measurements. **a)** Calculated residence times (*τ*_cal_) of opioids (blue bars) and their experimental residence times (gray bars). The *τ*_cal_ values (mean ± standard error of mean) were calculated from the bootstrapping analysis for 10,000 samples of size 15 each accounting for 15 simulated trajectories for each compound. **b)** Calculated vs. experimental residence times of 12 compounds (10 fentanyl and 2 morphine analogs). The black regression line was calculated using all 12 compounds. The purple line was calculated using only FEN and its its R4 and/or R1 modified derivatives. *r* represents the Pearson’s correlation coefficient. A logarithm scale is used. **c)** Measured dissociation constant(*K*_d,exp_) vs. calculated residence times of the 12 compounds. **d)** Calculated residence times vs. naloxone inhibitory constant (*K*_i,NLX_) of 9 of the compounds. *K*_i,NLX_ refers to the naloxone concentration that produces 50% reduction of the maximal agonist occupancy in mOR. The experimental values were obtained by Mann et al.^11^

Interestingly, with the Pearson’s *r* value of −0.87, the calculated *τ*’s has a stronger correlation with the experimental *K_d_*’s than the experimental *τ*’s (Fig. 2c). The improved correlation may be partly attributed to REMI, for which the calculated *τ* is larger than fentanyl, consistent with the experimental *K*_d_’s but not the experimental *τ*’s. Another contributor to the improved correlation is ALF, for which the calculation predicted that it has the smallest *τ* value among all compounds studied, consistent with the trend in the experimental *K*_d_ but not the experimental *τ*. In fact, the smallest experimental *τ* is given by NLX and not ALF.

Another property that is intuitively related to the residence time is *K*_i,NLX_, which is the concentration of NLX that produces half maximum inhibition of opioid binding. Interestingly, the calculated *τ*’s vs. *K*_i,NLX_’s gives a correlation coefficient of 0.87, which is identical to that for *τ*_calc_’s vs. *K*_d,expt_’s. Note, the experimental *K*_i,NLX_ values of REMI and ALF were not obtained (Table S1).

The experimental *τ* and *K*_d_ values of morphine derivatives studied here, MOP, NLX, NTX, and BUP, have been reported in two previous studies.^28,29^ The comparison of *τ*_exp_ and *K*_d_ values with *τ*_cal_ values gave *r* values greater than 0.9 (Fig. S3). Importantly, these studies showed that BUP has the longest residence times and smallest dissociation constant among the four opioids, in agreement with our calculations. BUP also gave a longer *τ*_exp_ than NLX in the experiment by our colleague Mann et al.,^11^ although the uncertainty for BUP is larger than NLX.

### Binding modes of fentanyls and morphinans

So far, all published X-ray structures of mOR, including those in complex with the morphinan agonist BU72,^16^ antagonist *β*-FNA,^24^ and endogenous peptide analog agonist DAMGO,^30^ feature a salt bridge between the charged amine and the conserved D147 on the transmembrane helix 3 (TM3). However, our recent study of FEN at mOR based on the enhanced sampling methods, WE and metadynamics simulations, revealed that, in addition to the D147 binding pose, fentanyl can rotate itself and insert deeper into mOR while donating a hydrogen bond (h-bond) to the *ϵ* nitrogen of the conserved H297 on TM6.^12,17^ The metadynamics simulations further suggested that the h-bond with H297 may prolong fentanyl’s residence time (*τ*), as without it (i.e., in the N_*ϵ*_ protonated or doubly protonated state), the *τ* value decreases by one or two orders of magnitude.^12^

In light of the above findings, we examined the interactions of the 19 fentanyl and morphine derivatives with D147 and H297 by calculating the approximate free energy surfaces (FES’) projected onto the distances between the ligand and D147 or H297 (Fig. 3 and SI). The presence of a salt bridge with D147 and/or hydrogen bond with H297 can be discerned from the local free energy minima (Fig. 3, highlighted by solid and dashed boxes). Interestingly, CAR, which has the longest experimental and calculated *τ*’s within the fentanyl analogs, forms both D147 and H297 interactions (Fig. 3a and d). By contrast, ALF, which has the shortest calculated *τ* and one of the shortest experimental *τ*’s, displays only the salt-bridge interaction with D147 (Fig. S4). Although this comparison is consistent with the hypothesis that H297 h-bond prolongs the residence time, SUF, which has a longer *τ* than fentanyl according to both experiment and simulation, does not interact with H297 (Fig. 3b and e). Even more unexpectedly, IBUF, FIBUF, and VAF form the h-bond with H297 rather than the salt bridge with D147 in simulations (Fig. 3c, f and Fig. S5).

**Figure 3.**
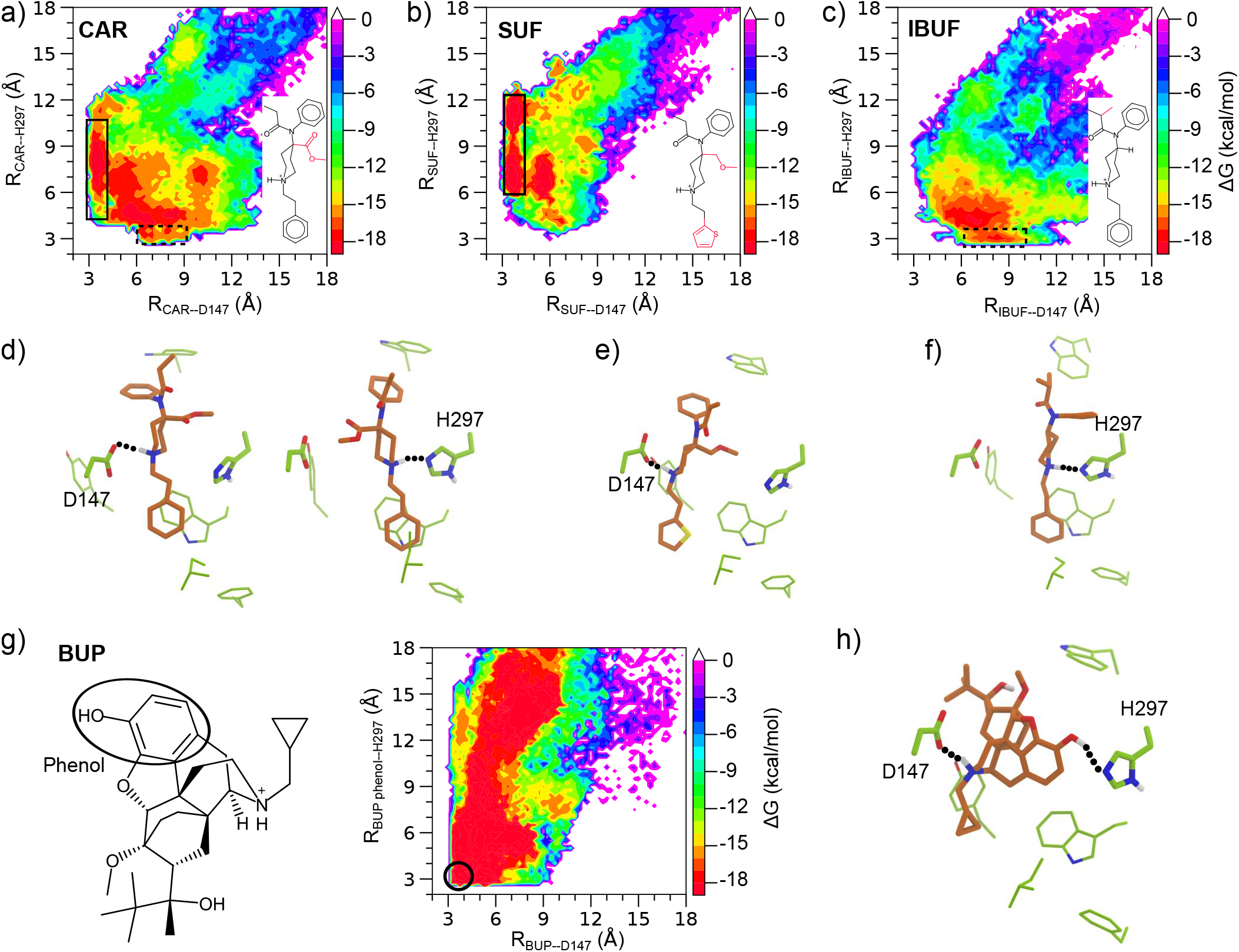
Binding modes of fentanyls and morphinans. **a, b, c)** Approximate free energy surfaces (FES’) of CAR, SUF and IBUF projected onto the distances between the piperidine nitrogen and D147 (carboxylate carbon) or H297 (nearest imidazole nitrogen). The FES regions representing the D147 or the H297 interactions are highlighted by solid and dashed boxes, respectively. **d)** Snapshots of the D147-(Left) and H297-bound (Right) poses of CAR. **e)** A snapshot of D147-bound pose of SUF. **f)** A snapshot of the H297 bound pose of IBUF. **g) Left.** Structure of BUP with the phenol ring highlighted. **Right.** Approximate FES of BUP projected onto two distances: between its amine and D147 (carboxylate carbon), and between the oxygen atom of the phenol (black circle) and H297 (nearest imidazole nitrogen). **h)** A snapshot of a bound pose of BUP where both the D147 and H297 interactions are formed. The unbiased free energies were calculated using the reweighting protocol ^26^ in PLUMED.^27^

Turning to the morphine derivatives, the charged amine group formed a stable salt bridge with D147 for all four compounds; however, none of them showed a h-bond between the charged amine and H297 (Fig. S6). Interestingly, BUP forms a salt bridge with D147 and a h-bond between the phenol hydroxyl group and H297 at the same time (Fig. 3g and h). The phenol-H297 h-bond was also observed in the simulations of NLX and NTX, albeit to a lesser extent as BUP (Fig. S7). Interestingly, the FES of BUP also shows a local minimum with the amine–D147 distance of 7–9 Å and phenol–H297 distance of 13–15 Å, representing a loosely bound state, which is consistent with a previous metadynamics study of BUP-mOR dissociation based on different collective variables. ^21^

### Residence time of fentanyl analogs is modulated by the substituent-mOR interaction energies

Since the fentanyl residence time is significantly affected by the R4 and/or R1 substitution (Fig. 2b and c), we hypothesized that they may be related to the interaction between the substituent and mOR. To test this hypothesis, we calculated the unbiased distributions of the interaction energies between R4/R1 and mOR obtained from the trajectories of CAR, LOF and REMI, METM, SUF and ALF and compared them with that of FEN. While R4 is a hydrogen for FEN, it is substituted by a methyl ester group in CAR, LOF and REMI and a methyl ether group in METM, SUF and ALF (Fig. 4a). Remarkably, the distributions of the R4-mOR interaction energies for these R4-substituted compounds are shifted by 5–6 kcal/mol to the negative values relative to fentanyl, indicating that a R4 methyl ester or methyl ether substitution stabilizes the interaction with mOR (Fig. 4b). The slightly lower energies for CAR/REMI/LOF relative to METM/SUF/ALF suggest that the methyl ester group forms a slightly more favorable interaction with mOR than the methyl ether group. CAR differs from FEN only in the R4 group, and the favorable interaction energy is consistent with the increased residence time and binding is consistent with the favorable interaction energy between the methyl ester and mOR, which can be attributed to a h-bond with the W318 (Fig. S9).

**Figure 4.**
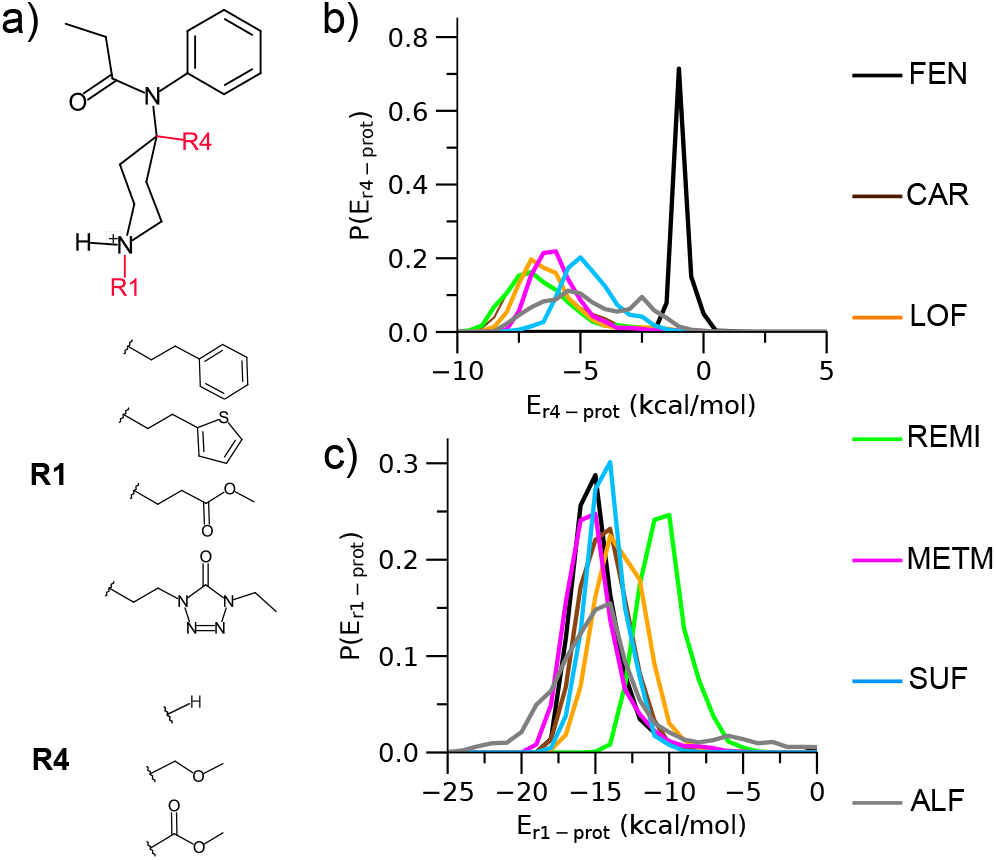
Residence time is modulated by the interactions between mOR and R4/R1 substituents. **a)** Chemical structure of the fentanyl scaffold highlighting the R4 and R1 substituents in the six compounds given on the right side. **b, c)** (Unbiased) distributions of the interaction energies between mOR and the R4 (b) or R1 (c) group in FEN (black), CAR (brown), LOF (orange), REMI (green), METM (magenta), SUF (cyan), and ALF (gray). The energies were calculated using the portions of metadynamics trajectories before the compound exited the protein (defined as *z* position relative to D147 greater than 15 Å as in our previous work^12^). The distributions were calculated using a reweighting protocol in PLUMED.^27^

Turning to the R1 group, CAR, LOF, METM have the same phenethyl group as FEN, which explains the similar distributions of the R1–mOR interaction energies (Fig 4c). SUF also shows a similar distribution, suggesting that the substitution of thiophene for phenethyl has a negligible effect (Fig 4c). By contrast, the distribution of REMI is right shifted by nearly 5 kcal/mol (Fig 4c, green curve), suggesting that the substitution of ethyl methyl ester for phenethyl is highly unfavorable, which may be attributed to the loss of the aromatic interaction of phenyl ring (in FEN) with W293.^12^ ALF shows the broadest distribution (Fig 4c, gray curve), suggesting that the interaction of ethyl-5-oxotetrazole and mOR is flexible, which is consistent with the very weak salt bridge between the piperidine amine and D147 (Fig. S4e). Indeed, the experimental *K_d_* value for ALF is the highest among all compounds studied.^11^

### Machine learning identifies the residue-substituent pairs important for fentanyl dissociation kinetics

The aforementioned analysis suggests that the mOR-R1/R4 interaction energies modulate the residence times of fentanyls; however, a more detailed understanding of the structure-kinetics relationship requires the consideration of a very large number of interactions between fentanyl substituents and mOR residues. To allow this, we enlisted a Machine Learning (ML) approach, in which tree-based regression models are trained to predict the *τ*_cal_ values using the interaction energies as features. Once accurate models are obtained, feature importance can be analyzed to identify important kinetics modulators.

The following ML workflow was implemented (Fig. 5a). First, for each ligand the trajectories were combined and randomly sampled followed by the calculation of the interaction energies between each substituent (R1, R2, or R4) and 272 mOR residues (terminal loop and a few other residues were excluded). To reduce noise and overfitting, the residue-substituent pairs were filtered based on the magnitude of interaction energies and their distributions for all compounds (reweighting was applied). Ultimately, 24 pairs survived (see Fig. S9–S11 for the reweighted distributions of the interaction energies). Next, we proceeded to the next round of random samplings and data augmentation followed by training the tree-based regression models to predict log_10_*τ*_cal_ values using PyCaret package^31^ (see Methods). The training and test sets were split in the 80:20 ratio and a 10-fold cross validation was used for model validation and hyperparameter tuning. The ML protocol (training, test, and feature importance calculation) was repeated 100 times to remove dataset bias. Among the six algorithms, Extra Trees, Random Forest, Gradient Boosting, and Extreme Gradient Boosting have similarly good performances, with the Pearson’s *r*^2^ value near or above 0.99 and root mean squared error below 0.1 (Table S2 and Fig. S12). Therefore, we continued with the first four models to rank features. Both the permutation based feature importance scores and SHAP (SHapley Additive exPlanations) values were calculated. The feature importance score represents the decrease in a model score (test) when a single feature value is randomly shuffled,^32^ while the SHAP value shows the impact of each individual feature value to each data point.^33^ To reduce overfitting, we removed 9 features with the lowest scores and used the top 15 features to retrain the four tree-based models (Table S3 and Fig. S13 and S14). The order of the feature importance scores and SHAP values from the four models are nearly identical to the corresponding ones using the larger feature set (Fig. S15– S18).

**Figure 5.**
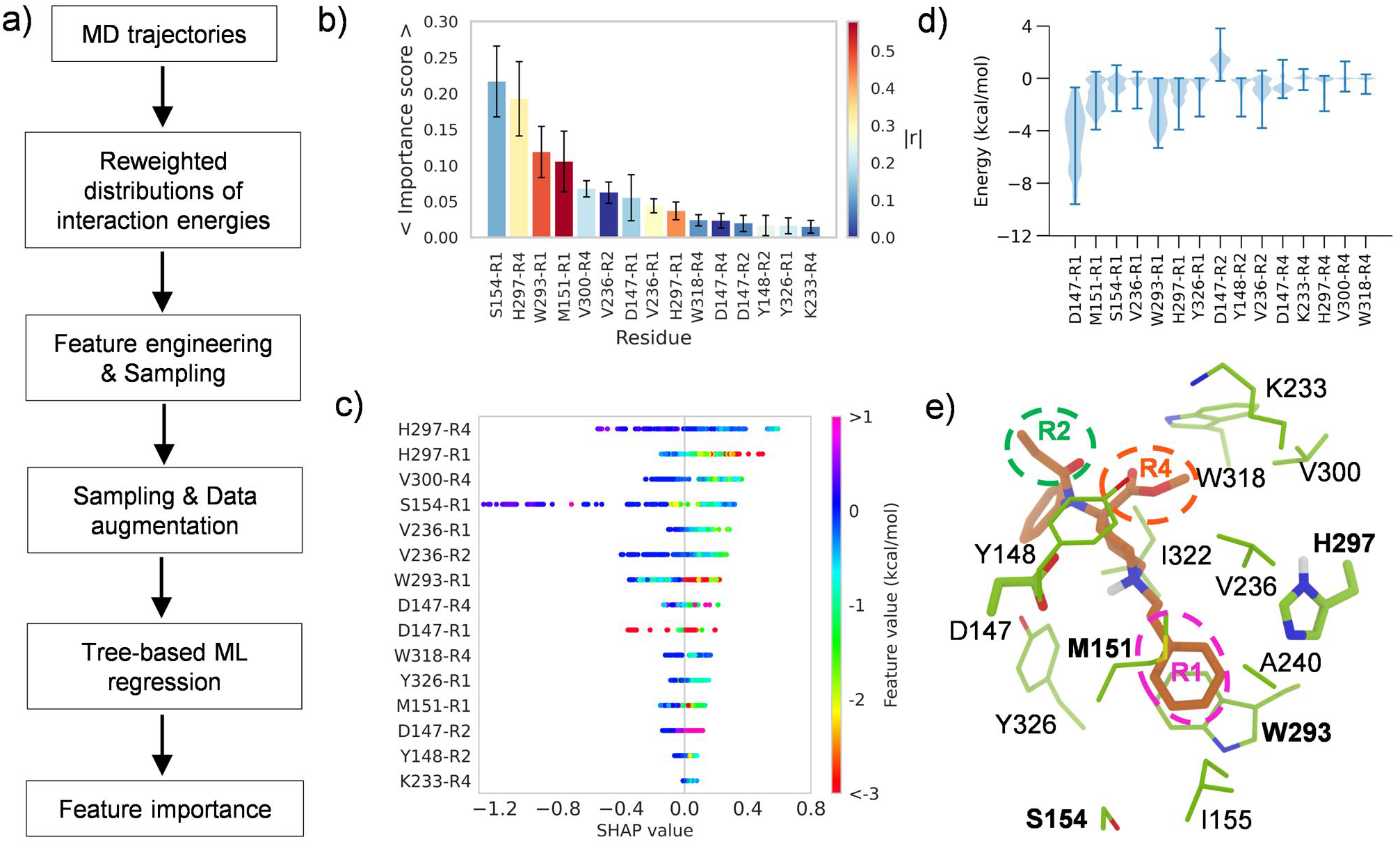
Machine learning (ML) identified structure modulators of dissociation kinetics. **a)** Steps in the ML analysis workflow. **b)** Feature importance scores calculated from the Random Forest models. The average and standard deviations (as error bars) were calculated over 100 trials. The color coding indicates the absolute value of the Pearson’s correlation coefficient for relating the interaction energies and the *τ* values (Fig. S13). **c)** A bee swarm plot of the SHAP values showing how individual features impact the predicted *τ* values from the random forest model. The features are ordered by the positive SHAP values. **d)** Violin plots of the reweighted interaction energies. The maximum, median, and minimum values are indicated by upper, middle and lower horizontal bars, respectively. **e)** A representative snapshot showing the contacts between CAR and nearby residues. The R1, R2 and R4 substituents are highlighted. Residues in bold font are involved in the most important features identified by the four ML models.

According to the average importance scores of the four models, the most important residue-substituent pairs for kinetics modulation include S154-R1, H297-R4, and W293-R1, and M151-R1 (Fig. 5b and Fig. S16). Note, while the H297-R4, and W293-R1, and M151-R1 interactions show at least some degree of correlation with the residence times (*r* value greater than 0.36), the S154-R1 interactions have a very low correlation with the residence times (*r* value of 0.2, Fig. 5b and Fig. S14). Interestingly, the SHAP value plots show that the residue-substituent pairs that have the largest positive impact on the residence times include H297-R4 and H297-R1 (Fig. 5c and Fig. S18). Closer examination of the bee swam plot shows that the stronger H297-R1/R4 interaction energies (more negative feature values) are associated with larger predicted residence times (positive SHAP value, Fig. 5c and Fig. S18). The prominent interaction between H297 and R1 is indeed

Trajectory analysis showed that the aromatic sidechains of H297 can form pi-pi stacking with the R1 phenyl ring of fentanyls (Fig. 5d), which may explain the increase in residence time. Conversely, loss of the pi-pi stacking, e.g., substitution of the phenyl with a non-aromatic group as in REMI, ALF, MN-FEN, and MNCAR significantly weakens the H297-R1 and W293-R1 interactions, which may explain the decrease in the residence time (Fig. 5e).

## CONCLUDING DISCUSSION

We applied a recently developed protocol^12^ based on well-tempered metadynamics simulations ^13,14^ to predict the residence times of fentanyls and morphinans at the mOR. Although the overall correlation between the calculated and most recent experimental *τ*’s^11^ is modest (Pearson’s *r* value of 0.61), exclusion of the highly similar R2 modified compounds (among them FBUF and FIBUF have large experimental errors) led to an increased *r* value of 0.83. Interestingly, even with the inclusion of the R2 modified compounds, the calculated *τ*’s correlate well with the experimental dissociation constants (*K_d_*),^11^ with *r* of −0.87. Notably, consistent with the largest experimental Kd value, ALF has the smallest calculated *τ*, which is 60 times smaller than NLX. However, the experimental *τ* of ALF is larger albeit on the same order of magnitude as NLX. Consistent with the second largest experimental Kd, REMI has the second shortest calculated *τ*, which is half of the *τ* for NLX; however, its experimental *τ* is nearly 20 times of the value for NLX. On the large end of the *τ* scale, both simulation and experiment ranked BUP and CAR as the top and second top compounds with the largest *τ*’s. As to the experimental *K_d_*’s, CAR and BUP have the first and second smallest values. Importantly, the calculated *τ*’s also correlated well with the experimental naloxone inhibitory constants, with the Pearson’s *r* of 0.87. Although the comparison does not include ALF and REMI, for which the experimental values are not available, this level of correlation suggests that computational prediction of *τ* values may be useful in the evaluation of naloxone dosing strategies for reversing opioid overdose. The underestimation of *τ* is consistent with previous metadynamics based kinetics calculations ^15^ and a recent calculation of morphine and buprenorphine based on infrequent metadynamics simulations.^21^ It may also reflect the sensitivity to the ligand force field parameters as demonstrated recently.^35^

In addition to the dissociation kinetics, the metadynamics simulations provided insights into the mechanism of opioid-mOR binding. Consistent with our previous work on fentanyl-mOR binding and dissociation,^12,17^ the present data of 15 fentanyls and 4 morphinans showed that while the majority of compounds, in particular morphinans, prefer to bind mOR via a salt bridge with D147, a few fentanyls, such as FUR, VAF, IBUF, FEN, FBUF, and FIBUF, can form either a salt bridge with D147 or a h-bond with H297. Surprisingly, among the latter ones, some fentanyls with the R2 modification, e.g., VAF and IBUF, prefer the binding mode via the h-bonding with H297. The precise binding mode of the fentanyls awaits experimental verification and further examination using unbiased MD simulations.

An important contribution of this work is the finding that the residence time change due to a substituent modification on the fentanyl scaffold is correlated with the substituent-mOR interaction energy. To systematically explore the interactions, we used the tree-based ML models trained to predict the fentanyls’ residence times to tease out the substituent-mOR pairs that are the most important modulators. Surprisingly, the ML models showed that the H297-R1/R4 interactions contribute the most to prolonging the residence time. This is in a stark contrast to the D147-R1 interaction, which is the strongest among all residue-substituent interactions but has little impact on the residence time.

The major caveat of the current work is the starting configurations used for the metadynamics simulations. As we were preparing the manuscript for submission, the cryo-EM structures of mOR in complex with fentanyl ^34^ and the closely-related lofentanil ^36^ were published, revealing that fentanyl is in a different orientation as the starting configuration used for simulations (Fig. 6a). Subsequently, we searched the metadynamics trajectories and found that similar configurations were captured by two out of the fifteen trajectories (Fig. 6b). The unbiased approximate free energy surface projected onto fentanyl’s orientation (defined using the piperidine ring angle relative to the membrane normal z axis) and z position or the piperidine-D147 salt bridge interaction distance shows that the minimum regions representing the cryo-EM like and the starting configurations are within 3 kcal/mol in free energy (Fig. 6d). Given the limited sampling, it is likely that the alternative configurations such as that captured by the cryo-EM model are undersampled leading to errors in the kinetics estimation. This is a major limitation of the current work. A second limitation of the work is the small number of experimental data points to compare with. Notwithstanding, our work provides a proof-of-concept approach for elucidating the structure-kinetics relationships of opioids. Unlike the conventional ligand-only ML models, the present approach combines ligand chemical structure with dynamical interactions between ligand and receptor. We envision that once accurate ML models are trained, predictions for new compounds can be rapidly made as only short MD or metadynamics simulations are needed to generate descriptors. Thus, we anticipate that ML models can be rapidly deployed to evaluate newly emerging synthetic opioids and inform overdose prevention strategies.

**Figure 6.**
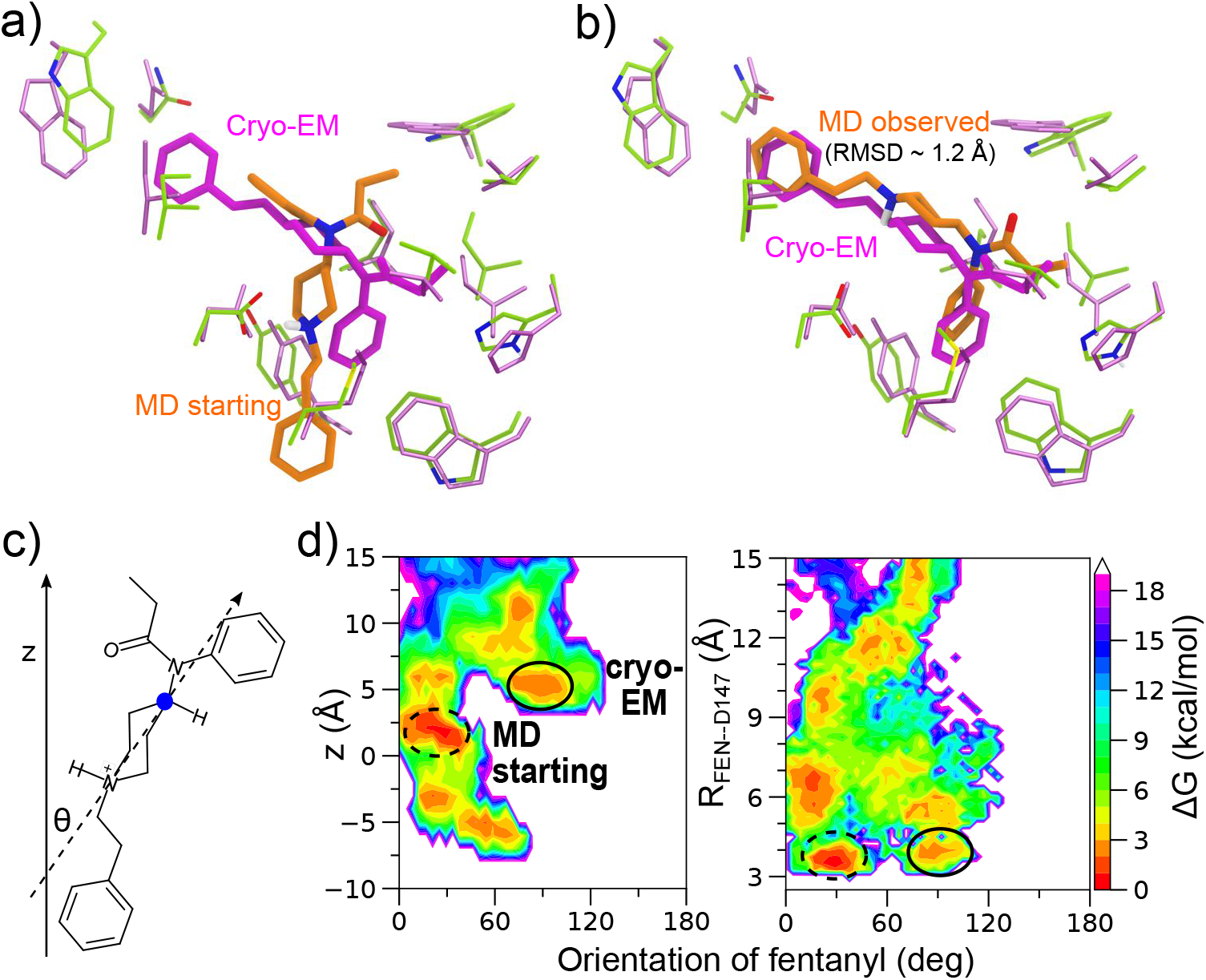
Metadynamics simulations captured fentanyl configurations resembling a newly determined cryo-EM model of human mOR in complex with fentanyl. **a,b)** Overlay of the fentanyl configuration (magenta) from the cryo-EM model of the mOR-fentanyl complex (PDB entry 8EF5,^34^ resolution 3.3 Å) onto the starting configuration (orange, **a**) of the metadynamics simulations and the most similar configuration (RMSD ~1.2 Å, **b**) from two of the metadynamics trajectories. **c)** Illustration of fentanyl’s orientation based on the piperidine angle (*θ*) relative to the normal vector z and the vector between the amine nitrogen and the opposite carbon atom (blue dot) in the piperidine group. **d)** Approximated free energy surface projected onto fentanyl’s orientational angle and z-position (left) or the piperidine–D147 distance (right). The latter distance is from the amine nitrogen to the nearest carboxylate oxygen. The regions representing the cryo-EM like and metadynamics starting configurations are highlighted by solid and dashed circles, respectively. The reweighted free energies were calculated from the two trajectories where the cryo-EM like configurations were observed.

## Supporting Information (SI)

Detailed Methods and protocols; Table S1 and S2 contain the summary of calculations and experimental data and assessment of ML model qualities. Figure S1 contains the structures of the opioids. Figure S2-S3 contain the additional comparisons between calculations and experiments. Figure S4-S8 contain additional analysis of trajectories. Figure S9-S15 contains additional data related to ML analysis.

## Data and Software Availability

The input files of all simulations as well as analysis and plotting scripts may be accessed at https://github.com/janashenlab/mOR_fentanyls. The molecular dynamics trajectories can be obtained from the authors upon request.

## Acknowledgement

We thank Pratyush Tiwary for the helpful advice on the residence time calculations from metadynamics simulations. We acknowledge the financial support from FDA’s Safety Research Interest Group and appointment to the Research Participation Programs at the Oak Ridge Institute for Science and Education through an interagency agreement between the Department of Energy and FDA. Financial support from the National Institutes of Health (R01GM098818) to J.S. is also acknowledged.

Parts of this study used the computational resources of the High performance Computing clusters at the FDA Center for Devices and Radiological Health, and the Extreme Science and Engineering Discovery Environment (XSEDE project number BIO210075) which is supported by National Science Foundation grant number ACI-1548562. Note, the findings and conclusions in this manuscript have not been formally disseminated by the FDA and should not be construed to represent any agency determination or policy.

## Dedication

J.S. would like to dedicate this work to a friend and other mothers who have lost their child to fentanyl overdose.

## References

(1) Ramirez, J.-M.; Burgraff, N. J.; Wei, A. D.; Baertsch, N. A.; Varga, A. G.; Baghdoyan, H. A.; Lydic, R.; Morris, K. F.; Bolser, D. C.; Levitt, E. S. Neuronal Mechanisms Underlying Opioid-Induced Respiratory Depression: Our Current Understanding. J Neurophysiol 2021, 125, 1899–1919.

(2) Vass, M.; Kooistra, A. J.; Yang, D.; Stevens, R. C.; Wang, M.-W.; de Graaf, C. Chemical Diversity in the G Protein-Coupled Receptor Superfamily. Trends in Pharmacological Sciences 2018, 39, 494–512.

(3) Burns, S. M.; Cunningham, C. W.; Mercer, S. L. DARK Classics in Chemical Neuroscience: Fentanyl. ACS Chem. Neurosci. 2018, 9, 2428–2437.

(4) Comer, S. D.; Cahill, C. M. Fentanyl: Receptor Pharmacology, Abuse Potential, and Implications for Treatment. Neurosci. Biobehav Rev. 2019, 106, 49–57.

(5) Vardanyan, R. S.; Hruby, V. J. Fentanyl-Related Compounds and Derivatives: Current Status and Future Prospects for Pharmaceutical Applications. Future medicinal chemistry 2014, 6, 385–412.

(6) Armenian, P.; Vo, K. T.; Barr-Walker, J.; Lynch, K. L. Fentanyl, Fentanyl Analogs and Novel Synthetic Opioids: A Comprehensive Review. Neuropharmacol. 2018, 134, 121–132.

(7) Ringuette, A. E.; Spock, M.; Lindsley, C. W.; Bender, A. M. DARK Classics in Chemical Neuroscience: Carfentanil. ACS Chem. Neurosci. 2020, 11, 3955–3967.

(8) Misailidi, N.; Papoutsis, I.; Nikolaou, P.; Dona, A.; Spiliopoulou, C.; Athanaselis, S. Fentanyls continue to replace heroin in the drug arena: the cases of ocfentanil and carfentanil. Forensic Toxicol. 2018, 36, 12–32.

(9) Skolnick, P. Treatment of overdose in the synthetic opioid era. Pharmacol. Thera 2022, 122, 108019.

(10) Moss, R. B.; Carlo, D. J. Higher Doses of Naloxone Are Needed in the Synthetic Opioid Era. Subst. Abuse Treat. Prev. Policy 2019, 14, 6.

(11) Mann, J. et al. Development of a Translational Model to Assess the Impact of Opioid Overdose and Naloxone Dosing on Respiratory Depression and Cardiac Arrest. Clin. Pharmacol. Therapeut. 2022, 112, 1020–1032.

(12) Mahinthichaichan, P.; Vo, Q. N.; Ellis, C. R.; Shen, J. Kinetics and Mechanism of Fentanyl Dissociation from the *μ*-Opioid Receptor. JACS Au 2021, 1, 2208–2215.

(13) Laio, A.; Parrinello, M. Escaping Free-Energy Minima. Proc. Natl. Acad. Sci. 2002, 99, 12562–12566.

(14) Barducci, A.; Bussi, G.; Parrinello, M. Well-Tempered Metadynamics: A Smoothly Converging and Tunable Free-Energy Method. Phys. Rev. Lett. 2008, 100, 020603.

(15) Tiwary, P; Limongelli, V.; Salvalaglio, M.; Parrinello, M. Kinetics of Protein–Ligand Unbinding: Predicting Pathways, Rates, and Rate-Limiting Steps. Proc. Natl. Acad. Sci. USA 2015, 112, E386–E391.

(16) Huang, W. et al. Structural Insights into μ-Opioid Receptor Activation. Nature 2015, 524, 315–321.

(17) Vo, Q. N.; Mahinthichaichan, P.; Shen, J.; Ellis, C. R. How μ-Opioid Receptor Recognizes Fentanyl. Nat Commun 2021, 12, 984.

(18) Zwier, M. C.; Adelman, J. L.; Kaus, J. W.; Pratt, A. J.; Wong, K. F.; Rego, N. B.; Suárez, E.; Lettieri, S.; Wang, D. W.; Grabe, M.; Zuckerman, D. M.; Chong, L. T. WESTPA: An Interoperable, Highly Scalable Software Package for Weighted Ensemble Simulation and Analysis. J. Chem. Theory Comput. 2015, 11, 800–809.

(19) Ballesteros, J. A.; Weinstein, H. In Methods in Neurosciences; Sealfon, S. C., Ed.; Receptor Molecular Biology; Academic Press, 1995; Vol. 25; pp 366–428.

(20) Casasnovas, R.; Limongelli, V.; Tiwary, P.; Carloni, P; Parrinello, M. Unbinding Kinetics of a P38 MAP Kinase Type II Inhibitor from Metadynamics Simulations. J. Am. Chem. Soc. 2017, 139, 4780–4788.

(21) Lamim Ribeiro, J. M.; Provasi, D.; Filizola, M. A Combination of Machine Learning and Infrequent Metadynamics to Efficiently Predict Kinetic Rates, Transition States, and Molecular Determinants of Drug Dissociation from G Protein-Coupled Receptors. J. Chem. Phys. 2020, 153, 124105.

(22) Callegari, D.; Lodola, A.; Pala, D.; Rivara, S.; Mor, M.; Rizzi, A.; Capelli, A. M. Metadynamics Simulations Distinguish Short-and Long-Residence-Time Inhibitors of Cyclin-Dependent Kinase 8. J. Chem. Inf. Model. 2017, 57, 159–169.

(23) Tiwary, P.; Parrinello, M. From Metadynamics to Dynamics. Phys. Rev. Lett. 2013, 111, 230602.

(24) Manglik, A.; Kruse, A. C.; Kobilka, T. S.; Thian, F. S.; Mathiesen, J. M.; Sunahara, R. K.; Pardo, L.; Weis, W. I.; Kobilka, B. K.; Granier, S. Crystal Structure of the *M*-Opioid Receptor Bound to a Morphinan Antagonist. Nature 2012, 485, 321–326.

(25) Salvalaglio, M.; Tiwary, P.; Parrinello, M. Assessing the Reliability of the Dynamics Reconstructed from Metadynamics. J. Chem. Theory Comput. 2014, 10, 1420–1425.

(26) Bonomi, M.; Barducci, A.; Parrinello, M. Reconstructing the Equilibrium Boltzmann Distribution from Well-Tempered Metadynamics. J. Comput. Chem. 2009, 30, 1615–1621.

(27) Tribello, G. A.; Bonomi, M.; Branduardi, D.; Camilloni, C.; Bussi, G. PLUMED 2: New Feathers for an Old Bird. Comput. Phys. Commun. 2014, 185, 604–613.

(28) Cassel, J. A.; Daubert, J. D.; DeHaven, R. N. [3H]Alvimopan Binding to the μ Opioid Receptor: Comparative Binding Kinetics of Opioid Antagonists. Eur. J. Pharmacol. 2005, 520, 29–36.

(29) Pedersen, M. F.; Wróbel, T. M.; Märcher-Rørsted, E.; Pedersen, D. S.; Møller, T. C.; Gabriele, F.; Pedersen, H.; Matosiuk, D.; Foster, S. R.; Bouvier, M.; Bräuner-Osborne, H. Biased Agonism of Clinically Approved μ-Opioid Receptor Agonists and TRV130 Is Not Controlled by Binding and Signaling Kinetics. Neuropharmacol. 2020, 166, 107718.

(30) Koehl, A. et al. Structure of the μ-Opioid Receptor-Gi Protein Complex. Nature 2018, 558, 547–552.

(31) Ali, M. PyCaret: An open source, low-code machine learning library in Python. 2020; PyCaret version 1.0.

(32) Breiman, L. Random Forests. Mach. Learn. 2001, 45, 5–32.

(33) Lundberg, S. M.; Lee, S.-I. A Unified Approach to Interpreting Model Predictions. Advances in Neural Information Processing Systems. 2017.

(34) Zhuang, Y. et al. Molecular Recognition of Morphine and Fentanyl by the Human μ-Opioid Receptor. Cell 2022, 185, 4361–4375.e19.

(35) Capelli, R.; Lyu, W.; Bolnykh, V.; Meloni, S.; Olsen, J. M. H.; Rothlisberger, U.; Parrinello, M.; Carloni, P. Accuracy of Molecular Simulation-Based Predictions of *k* _off_ Values: A Metadynamics Study. J. Phys. Chem. Lett. 2020, 11, 6373–6381.

(36) Qu, Q. et al. Insights into Distinct Signaling Profiles of the μOR Activated by Diverse Agonists. Nat. Chem. Biol. 2022,

