## Supporting Information for "Structure-Kinetics Relationships of Opioids from Metadynamics and Machine Learning"

† *Division of Applied Regulatory Science, Office of Clinical Pharmacology, Center for Drug Evaluation and Research, United States Food and Drug Administration, Silver Spring, MD 20993, United States*

### List of Tables


S1 Calculated residence times in comparison to the related experimental data S12

S2 Performance comparison of tree-based regression models (Top 15 features) S13

S3 Performance comparison of tree-based regression models (all 24 features) S14

#### List of Figures

S1 Chemical structures of the simulated opioids . . . . . . . . . . . . . . . . . . . S15
S2 The calculated and experimental residence times of fentanyls with R4 and/or R1 modification correlate well . . . . . . . . . . . . . . . . . . . . . . . . . S16
S3 Correlation between the calculated residence times of the morphinans with the previous experimental measurements . . . . . . . . . . . . . . . . . . . . S17
S4 Interactions between fentanyls and the mOR residues . . . . . . . . . . . . S18
S5 Approximate free energy surfaces illustrating the D147 salt bridge and the H297 hydrogen bond with the rest of fentanyls with R4 and/or R1 modification S19
S6 Approximate free energy surfaces illustrating the D147 salt bridge and the H297 hydrogen bond with the rest of fentanyls with R2 modification . . . . . S20
S7 Piperidine-D147 and piperidine-H297 interactions for morphinans . . . . . . S21
S8 Amine-D147 and phenol-H297 h-bond interactions for the morphinans . . . S22
S9 Interaction energies between selected mOR residues and the R1 group . . S23
S10 Interaction energies between selected mOR residues and the R2 group . . S24
S11 Interaction energies between selected mOR residues and the R4 group . . S25
S12 Model qualities of the tree-based regression models (all 24 features) . . . . S26
S13 Model qualities of the tree-based regression models (15 features) . . . . . . S27
S14 Correlation between the interaction energies and the calculated residence times . . . . . . . . . . . . . . . . . . . . . . . . . . . . . . . . . . . . . . . S28
S15 Average feature importance scores for the top four models (all 24 features) S29
S16 Average feature importance scores for the top four models (top 15 features) S30
S17 Beeswarm SHAP value plots for the top four models (all 24 features) . . . . S31
S18 Beeswarm SHAP value plots for the top four models (top 15 features) . . . S32

### Methods and protocols

**Ligand similarity matrix calculations.** The structural similarities of the opioids were evaluated using Tanimoto coefficients ($T_c$)[S1] calculated by the RDKit package[S2] We first converted the SMILES (Simplified molecular-input line-entry system) strings of molecules to the 1,024-bit Extended-Connectivity fingerprints with the radius of 2 bonds.[S3] Once the fingerprints were obtained, $T_c$ between two molecules was calculated as[S4] $Tc(A, B) = c/(a + b - c)$, where $a$ and $b$ represent the number of features present in molecule A, B, respectively, while $c$ represents the number of features common to A and B.

**System preparation.** 15 fentanyl (FEN, BUF, IBUF, FBUF, FIBUF, FUR, VAF, CAR, LOF, REMI, METM, SUF, ALF, MNCAR, MNFEN) and 4 morphine analogs (BUP, NLX, NTX, and MOP) were studied in this work. As in our previous metadynamics study of FEN-bound mOR,[S5] the starting structures of mOR in complex with the fentanyl analogs were modified from a representative snapshot taken from the global free energy minimum sampled by the WE simulations of FEN-bound mOR.[S6] The latter simulations were initiated from the MD relaxed docked structure generated using the X-ray structure of the agonist BU72-bound mOR (PDB entry 5C1M)[S7] as a template and root-mean-square deviation (RMSD) from it as a progress variable. The same X-ray structure was used as a template to dock MOP and BUP into mOR. For docking NLX and NTX into mOR, the antagonist $\beta$-FNA bound X-ray crystal structure of mOR (PDB entry 4DKL)[S8] was used as a template. Although the crystal structures are of mouse's mOR, the amino acid sequences of the human and mouse receptors are nearly identical. The protonation states of mOR were taken from our previous work, which were determined by the membrane-enabled hybrid-solvent continuous constant pH molecular dynamics (CpHMD) method with the pH replica-exchange protocol.[S9–S11] It was found that D114 ($D^{2.50}$) is charged and H297 ($H297^{6.52}$) samples the neutral Hid and Hie tautomer states at physiological pH. Since the previous metadynamics simulations of FEN-mOR dissociation[S5] found that the Hid state gave one order of magnitude slower kinetics than the Hie state, resulting in a better agreement with experiment, we adopted it in the current simulations. The rest of the titratable sites were found to be in the standard protonation states (Asp/Glu is charged, His is in the neutral Hie state, Lys is charged, and Cys is neutral) and were adopted as such. All fentanyls and morphinans carry a charge of $+1$. Following the previous protocols,[S5,S6] each of the aforementioned ligand-mOR complex structures was embedded in the POPC (1-palmitoyl-2-oleoyl-glycero-3-phosphocholine) lipid bilayer and solvated with water and $Na^+$/$Cl^-$ ions. The resulting system underwent stepwise MD equilibration and finally unrestrained MD to relax the conformation for at least 200 ns prior to the metadynamics simulations (details see our previous work[S5,S6]).

**Molecular dynamics protocol.** All simulations were performed with NAMD2.[S12] The protein and lipids were represented by the CHARMM36 protein and lipid force fields, respectively.[S13,S14] The CHARMM modified TIP3P model was used to represent water.[S15] The fentanyls and morphinans were represented by the CGenFF force field (version 3.0.1) obtained through the Paramchem server.[S16,S17] All bond and angles involving hydrogen atoms were constrained using the SHAKE algorithm[S18] to allow an integration timestep of 2 fs. The simulations were performed under periodic boundary and constant NPT conditions in a flexible cell with a constant xy ratio. The temperature was maintained at 310 K by Langevin dynamics with a damping coefficient $\gamma$ of 1 ps$^{-1}$, and the pressure was controlled at 1 atm by the Nosé-Hoover Langevin piston method.[S19,S20] The van der Waals interactions were smoothly switch off from 10 to 12 Å using a switching function. The particle mesh Ewald (PME) method[S21] was used to calculate long-range electrostatic energies with a sixth-order interpolation and a grid spacing of 1 Å. Most of the analyses were performed using VMD[S22] and PLUMED.[S23]

**Metadynamics simulations of ligand-mOR dissociation.** Metadynamics simulations were performed using the `Colvars` module[S24] in `NAMD2`,[S12] which implements the well-tempered metadynamics method.[S25,S26] The ligand-mOR dissociation process was accelerated through the deposition of a time-dependent Gaussian potential along two collective variables CV1 and CV2. Following our previous work,[S5] the CV1 was the ligand-mOR contact number, and CV2 was the ligand center-of-mass (COM) z position relative to that of the $C_\alpha$ atoms in the putative binding pocket according to the X-ray structure of BU72-bound mOR.[S7] These CVs do not presume a particular unbinding pathway. The residues in the binding pocket include Y75, Q124, N127, W133, L144, D147, Y148, M151, F152, L232, K233, V236, A240, W293, I296, H297, V300, W318, H319, I322, and Y326. The ligand-mOR contact number (CN) is defined using a differentiable switching function

$$\mathrm{CN} = \sum_{i \in \text{mOR}} \sum_{j \in \text{ligand}} \frac{1-(d_{ij}/R_{\text{cut}})^8}{1-(d_{ij}/R_{\text{cut}})^{16}}, \tag{S1}$$

where $d_{ij}$ is the distance between the heavy atom $i$ in ligand and $j$ in mOR and $R_{\text{cut}}$ is the effect cutoff distance which was set to 4.5 Å. Following the previous ligand-protein dissociation simulations,[S5] 15 independent metadynamics simulations with different random velocity seeds were carried out for each ligand. A cutoff of 15 Å for CV2 (z position) was used to define the unbound state. The robustness of this cutoff in range of 10–20 Å was examined previously.[S5] There were a total of 285 trajectories, with the aggregate sampling time of $\sim$15 $\mu$s. The Gaussian weight parameter $W$ (hill height) was set to 0.5 kcal/mol and the width $\sigma$ was set to 5 for CV1 and 0.5 Å for CV2. The tempering parameter $\Delta$T was set to 14T, where T is the simulation temperature 310 K. The PLUMED program[S23] was used to calculate the unbiased free energy profiles and the unbiased dissociation times (see below).

**Estimation of ligand-mOR residence times.** Following Casanovas[S27] and others,[S28] we first converted the individual metadynamics exit time $t_{\text{exit,meta}}$ to the real time $t_{\text{real}}$ by summing up the time steps rescaled at each step :

$$t_{\mathrm{real}} = \sum_{i=1}^{N} \delta t e^{\beta V_{\mathrm{meta}}(t_i)}, \tag{S2}$$

where $i$ is the step number for bias deposition, $N$ is the total number of bias deposition steps, $\delta t$ is the time step for bias deposition and $V_{\mathrm{meta}}(t_i)$ is the bias potential at time $t_i$. Following Salvalaglio et al.,[S29] the dissociation time ($\tau$) was estimated by fitting the empirical cumulative distribution (CDF) function which represents the probability of observing at least one dissociation event by time $t$, to the theoretical CDF for a Poisson process,

$$P_{n\geq 1} = 1 - e^{-t_{\mathrm{real}}/\tau}, \tag{S3}$$

where $\tau$ is the estimated residence time. The Kolmogorov-Smirnov (KS) test was used to test the null hypothesis that the sample of dissociation times extracted from metadynamics and a large sample of times randomly generated according to the theoretical probability density reflect the same underlying distribution.[S29] The null hypothesis is rejected if $p$-value < $\alpha$, where $\alpha$ is typically chosen as 0.05. To estimate the errors of $\tau$, we performed bootstrapping analysis with 10,000 samples. Following Palacio-Rodriguez et al,[S30] we used the samples that passed the KS test, defined by p-value $\geq$0.05, for calculation of the mean and standard error (SE) of $\tau$.

**Calculations of protein-ligand or residue-substituent interaction energies.** The interaction energy calculations were performed using the in-house tcl scripts for VMD.[S22] The interaction energy was defined as the sum of electrostatic and van der Waals energies between the protein (or an individual residue) and ligand (or a substituent group). All atoms and a dielectric constant of 4 were used in the calculations. For each ligand, the 15 trajectories (saved every 50 ps), with the initial 5 ns and the dissociated portion (CV2 greater than 15 Å, see above) discarded, were combined, resulting in 7,500–16,000 frames.

**Feature engineering.** For ligand, the aforementioned trajectory frames were randomly sampled 100 times and each gave 2,000 snapshots. For these snapshots, the interaction energies between the ligand and 272 mOR residues (excluding the terminal residues 52–64 and 340–347) were calculated and the average values were saved, which resulted in 272 residue-ligand interaction energies for each sample. To avoid noise-related artifacts and model overfitting, the following steps were performed to prune features. First, a mOR residue was removed if the absolute residue-ligand interaction energy is below 0.5 kcal/mol for all ligands. This led to 37 mOR residues (A113, D114, A117, Q124, I144, I146, D147, Y148, N150, M151, F152, S154, I155, L158, D216, E229, L232, K233, V236, F237, A240, F241, F289, I296, W293, H297, V300, K303, W318, C321, G325, I322, Y326, N328, S329, and N322) for further consideration. Next, the probability distributions of the residue-ligand interaction energies were examined. A residue was removed if it has weak interactions with all ligands, i.e., a narrow distribution (range is within $\sim$2 kcal/mol) and a peak near 0 kcal/mol. This step resulted in 16 residues: D147, Y148, N150, M151, S154, I155, K233, V236, A240, W293, I296, H297, V300, W318, I322 and Y326. For the above 16 residues and the fentanyl substituent R1, R2, and R4, the distributions of the residue-substituent interaction energies based on the combined trajectories with 37,500–80,000 frames were calculated (Fig. S8-10). In total, there were 720 distributions (16 residues $\times$3 R1/R2/R4 $\times$15 ligands) representing 48 residue-substituent contact pairs for each fentanyl analog. Note, due to the lack of data, R3 and R5 were not considered. The residue-substituent contacts with weak interactions, i.e., peak around $\sim$0 kcal/mol and range never extending below -2 kcal/mol for all ligands, were discarded. This allowed us to extract a total of 24 residue-substituent pairs (D147-R1, M151-R1, S154-R1, I155-R1, V236-R1, A240-R1, W293-R1, H297-R1, W318-R1, I322-R1, Y326-R1, D147-R2, Y148-R2, K233-R2, V236-R2, V300-R2, W318-R2, D147-R4, Y148-R4, K233-R4, H297-R4,

V300-R4, W318-R4, and I322-R4) as features for a supervised ML analysis.

**Machine learning analysis of kinetics modulators.** For each ligand, 2000 frames were randomly sampled from the 37,500–80,000 combined trajectory frames. From these frames the interaction energies of the 24 residue-substituent pairs were calculated and the average energies obtained from the reweighted distributions using the reweighting protocol in PLUMED (version 2.5)[S23] were saved. This resulted in 24 average energies (features) for each ligand. Since we only have 15 residence times (for 15 ligands), a data augmentation "trick" was applied in which we repeat the above protocol 100 times, resulting in 2,400 (24×100) energies for 1500 ligands. At this stage, there are more parameters but we will address this issue later.

The above energies were used as features to train tree-based regression models for predicting the $\log_{10}$ transformed calculated residence times $\log_{10}(\tau_{\rm cal})$ of fentanyls. The ML was performed using `PyCaret`[S31] which implements six popular tree-based regression methods, Extra Trees, Random Forest, Decision Tree, Adaptive Boosting, Gradient Boosting, and Extreme Gradient Boosting (Table S3). The training and test sets were split in a 80:20 ratio and a 10-fold cross validation was used for model validation and hyperparameter tuning. After tuning the hyperparameters using random grid search for 1000 steps, feature importance scores were calculated. To reduce possible dataset bias, we ran the ML procedure (training, testing, and feature importance calculation) for a total of 100 trials; in each trial, the splitting for training and test sets was random and independent. Among those tested models (Table S3 and Fig. S12), the Extra Trees, Random Forest, Gradient Boosting, and Extreme Gradient Boosting gave similarly good performances based on multiple evaluation metrics, including the mean average error (MAE), root mean squared error (RMSE), coefficient of determination ($R^2$), and mean absolute percentage error (MAPE); it gave the second best score for the root mean squared logarithmic error (RMSLE). Thus, we calculated the importance scores and SHAP values of the features for these four types of models.

#### Supplemental tables

**Table S1:** A summary of the calculated residence times of opioids in comparison to the measured residence times, dissociation constants, and naloxone inhibitory constants

| No. | Name | $\tau_{\rm cal}$ (sec) | Success bootstrapping samples | p-value for $\tau_{\rm cal}$ | $\tau_{\rm exp}$ (sec) | $K_{\rm d}$ (nM) | $K_{\rm i,NLX}$ (nM) |
|---|---|---|---|---|---|---|---|
| | | | | **Fentanyl analogs** | | | |
| Mainly R2 modified | | | | | | | |
| 1 | FEN | 89.1±1.9 | 3631 | 0.13 | 231±17 | 0.68±0.08 | 3.0±0.5 |
| 3 | BUF | 64.1±0.4 | 8255 | 0.23 | 188±26 | 0.76±0.16 | 1.6±0.3 |
| 4 | FBUF | 38.2±1.5 | 4306 | 0.15 | 65±26 | 0.71±0.26 | 1.7±0.5 |
| 7 | FIBUF | 26.8±0.3 | 5559 | 0.17 | 159±45 | 0.69±0.19 | 1.0±0.2 |
| 6 | IBUF | 52.9±0.4 | 8591 | 0.25 | 32±3 | 0.71±0.01 | 1.3±0.2 |
| 8 | FUR | 147.6±13.6 | 4248 | 0.11 | 184±54 | 0.31±0.04 | 3.3±0.5 |
| 5 | VAF | 151.7±1.9 | 4601 | 0.15 | - | - | - |
| R1 or/and R4 modified | | | | | | | |
| 15 | ALF | 0.35±0.004 | 8518 | 0.26 | 71±19 | 11.6±2.0 | - |
| 12 | REMI | 4.64±0.13 | 5114 | 0.12 | 481±59 | 1.45±0.10 | - |
| 10 | CAR | 367.8±3.9 | 5829 | 0.18 | 4049±199 | 0.05±0.04 | 4.1±0.6 |
| 14 | SUF | 141.7±1.3 | 7751 | 0.17 | 935±69 | 0.17±0.03 | 6.1±0.3 |
| 11 | LOF | 698.1±7.8 | 7258 | 0.16 | - | - | - |
| 2 | MNFEN | 0.42±0.003 | 8005 | 0.31 | - | - | - |
| 9 | MNCAR | 22.9±0.5 | 4410 | 0.12 | - | - | - |
| 13 | METM | 177.2±3.9 | 3894 | 0.14 | - | - | - |
| | | | | **Morphine analogs** | | | |
| 1 | BUP | 1652.3±12.4 | 5300 | 0.17 | 7692±1395 | 0.19±0.05 | 8.6±1.8 |
| 2 | MOP | 0.43±0.005 | 8838 | 0.28 | - | - | - |
| 3 | NLX | 6.22±0.16 | 3559 | 0.13 | 25±6 | 0.71±0.08 | - |
| 4 | NTX | 6.68±0.05 | 9480 | 0.34 | - | - | - |

Success samples are the number of 10,000 bootstrapping samples that passed the KS test defined by a p-value of ≥0.05. Following Palacio-Rodriguez et al,[S30] The mean and standard error of the mean (SEM) values of $\tau_{\rm cal}$ were estimated from the success samples. The p-value refers to the average p-value calculated from the success samples. $\tau_{\rm exp}$, $K_{\rm d}$ and $K_{\rm i,NLX}$ values were obtained by Mann et al.[S32] While the SE values of $K_{\rm d}$ and $K_{\rm i,NLX}$ for each of the compounds was reported, the one of its $\tau_{\rm exp}$ was estimated from the reported 95% confidence intervals,[S32] and is defined as (upper limit - lower limit)/3.92.

**Table S2:** Performance comparison of the regression models built with different tree-based machine learning (ML) methods (Top 15 features)

| Method | MAE | RMSE | $R^2$ | RMSLE | MAPE |
|---|---|---|---|---|---|
| Extra Trees | 0.022 | 0.055 | 0.996 | 0.024 | 0.017 |
| Random Forest | 0.029 | 0.087 | 0.991 | 0.037 | 0.024 |
| Gradient Boosting | 0.051 | 0.098 | 0.988 | 0.041 | 0.043 |
| Extreme Gradient Boosting | 0.044 | 0.098 | 0.988 | 0.040 | 0.035 |
| Decision Tree | 0.026 | 0.149 | 0.971 | 0.051 | 0.020 |
| Adaptive Boosting | 0.146 | 0.191 | 0.958 | 0.074 | 0.111 |

The model quality metrics listed are mean average error (MAE), root mean squared error (RMSE), coefficient of determination ($R^2$), root mean squared logarithmic error (RMSLE), and mean absolute percentage error (MAPE). These values were obtained based on 100 trials through `PyCaret`.[S31] The first four methods are similar in performance although the extra trees method is slightly better. The 15 most important features of the 24 features (D147-R1, M151-R1, S154-R1, V236-R1, H297-R1, W293-R1, Y326-R1, D147-R2, Y148-R2, V236-R2, D147-R4, K233-R4, H297-R4, V300-R4) identified by Extra Trees were used in the evaluation.

**Table S3:** Performance comparison of the regression models built with different tree-based machine learning (ML) methods (all 24 features)

| Method | MAE | RMSE | $R^2$ | RMSLE | MAPE |
|---|---|---|---|---|---|
| Extra Trees | 0.021 | 0.051 | 0.997 | 0.024 | 0.018 |
| Random Forest | 0.031 | 0.087 | 0.991 | 0.038 | 0.026 |
| Gradient Boosting | 0.051 | 0.098 | 0.989 | 0.041 | 0.044 |
| Extreme Gradient Boosting | 0.044 | 0.099 | 0.988 | 0.042 | 0.035 |
| Decision Tree | 0.025 | 0.140 | 0.974 | 0.048 | 0.019 |
| Adaptive Boosting | 0.144 | 0.187 | 0.960 | 0.073 | 0.109 |

The model quality metrics listed are mean average error (MAE), root mean squared error (RMSE), coefficient of determination ($R^2$), root mean squared logarithmic error (RMSLE), and mean absolute percentage error (MAPE). These values were obtained based on 100 trials through `PyCaret`.[S31] The first four methods are similar in performance although the extra trees method is slightly better.

### Supplemental figures

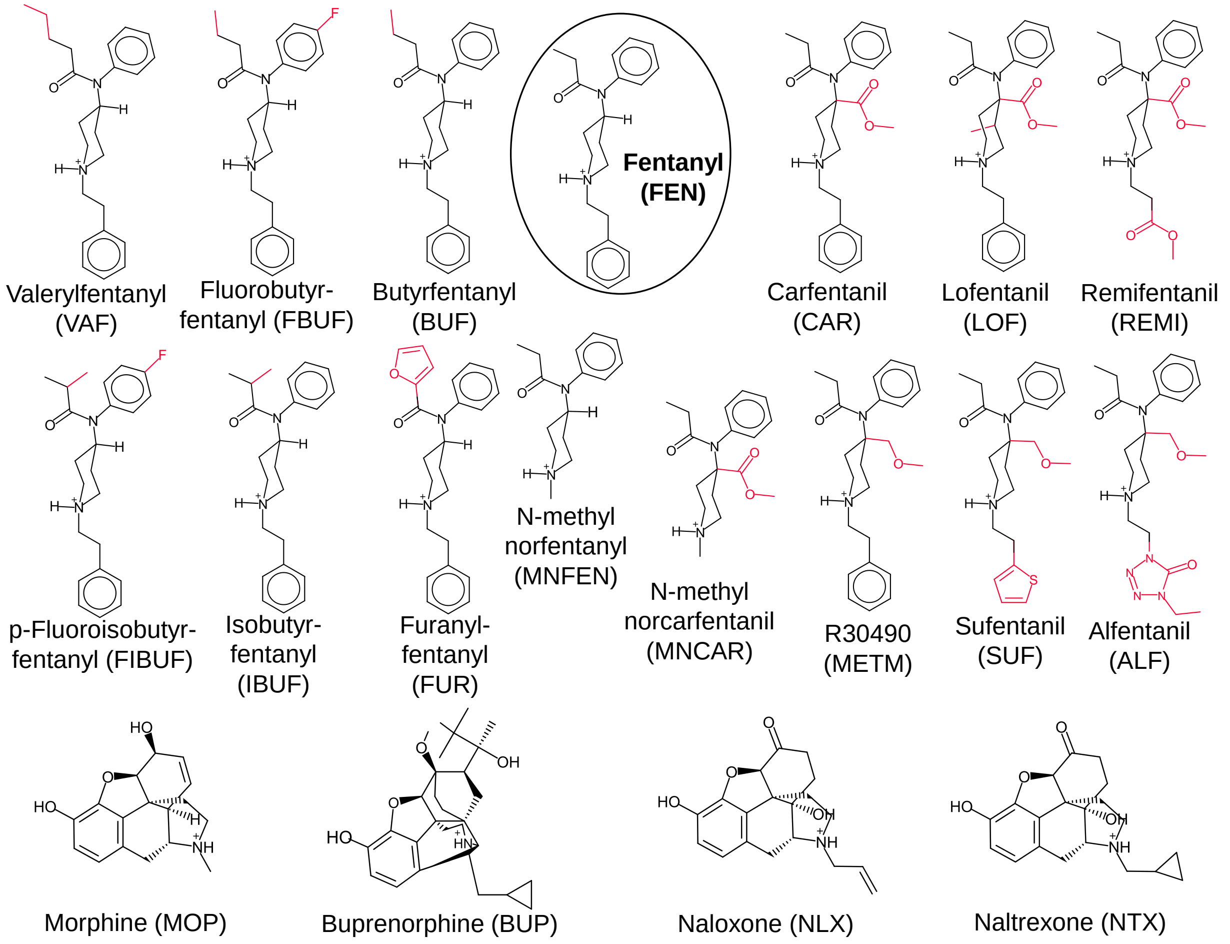


**Figure S1:** Chemical structures of the simulated opioids. Except MNFEN, the substituents in the fentanyl (FEN) analogs are highlighted in red.

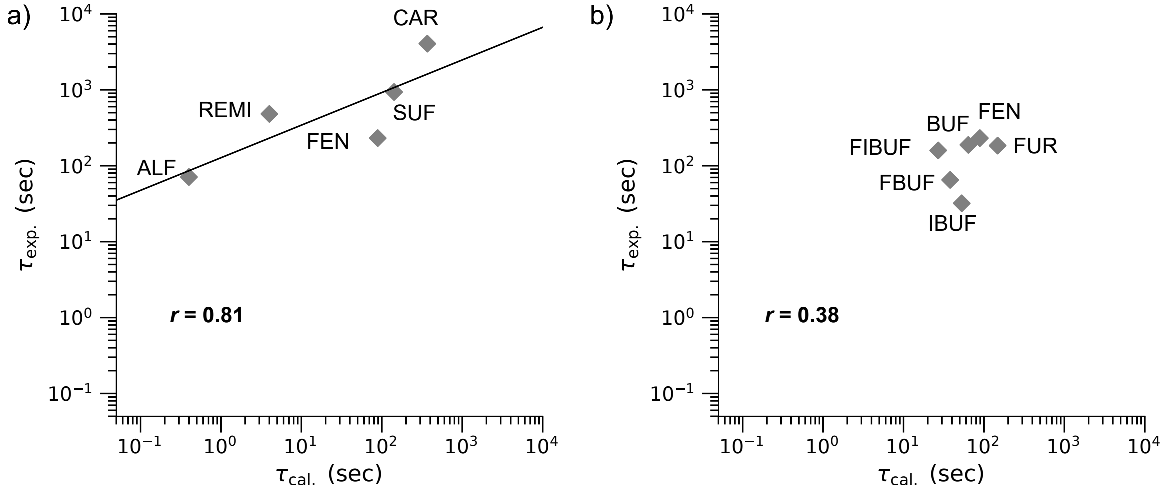


**Figure S2:** Correlation between the calculated and experimental residence times of fentanyls with R4 and/or R1 modification (a) or with R2 modification (b). $r$ is the Pearson's correlation coefficient. A logarithm scale is used. The experimental residence times were obtained by Mann et al[S32]

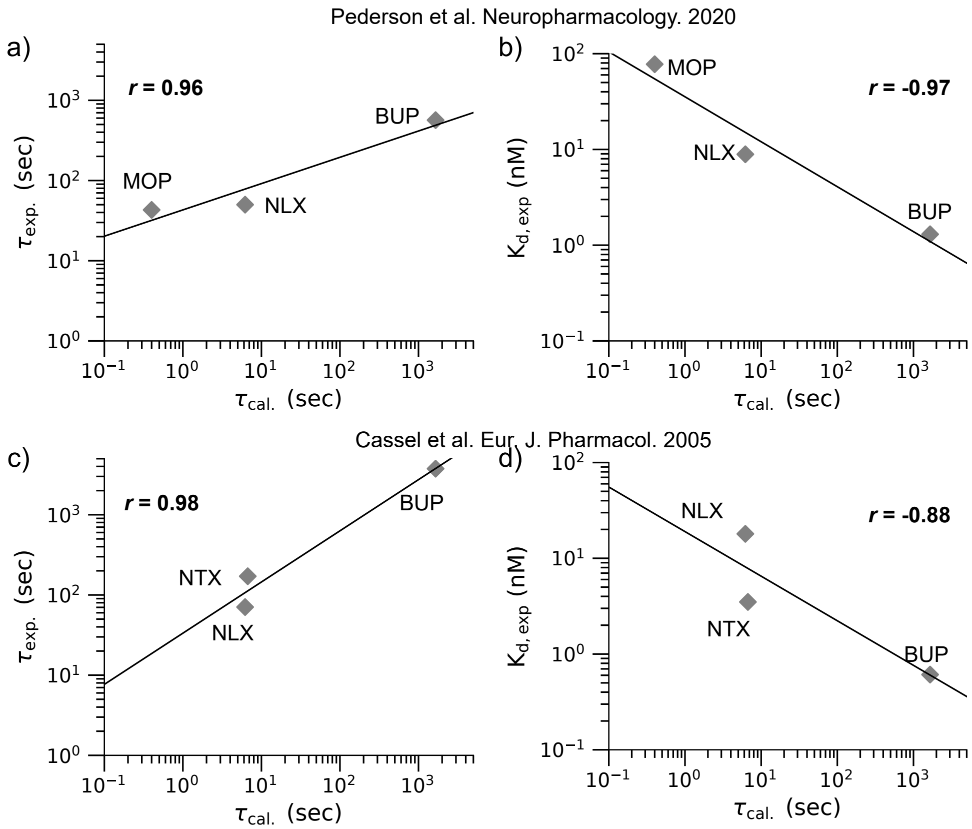


**Figure S3: Correlation between the calculated residence times of the morphinans with the previous experimental measurements. a–b)** Correlation between $\tau_{\rm cal}$ and $\tau_{\rm exp}$ or $K_{\rm d}$ of BUP, NLX and MOP determined by Peterson et al.[S33] The $\tau_{\rm exp}$ values for BUP, NLX and MOP are 566, 50 and 43 s, respectively.[S33] **c–d)** Correlation between $\tau_{\rm cal}$ and $\tau_{\rm exp}$ or $K_{\rm d}$ of BUP, NLX and NTX determined by Cassel et al.[S34] The $\tau_{\rm exp}$ values for BUP, NLX and NTX are 3,750, 71 and 171 s, respectively[S34] $r$ is the Pearson's correlation coefficient. A logarithm scale is used.

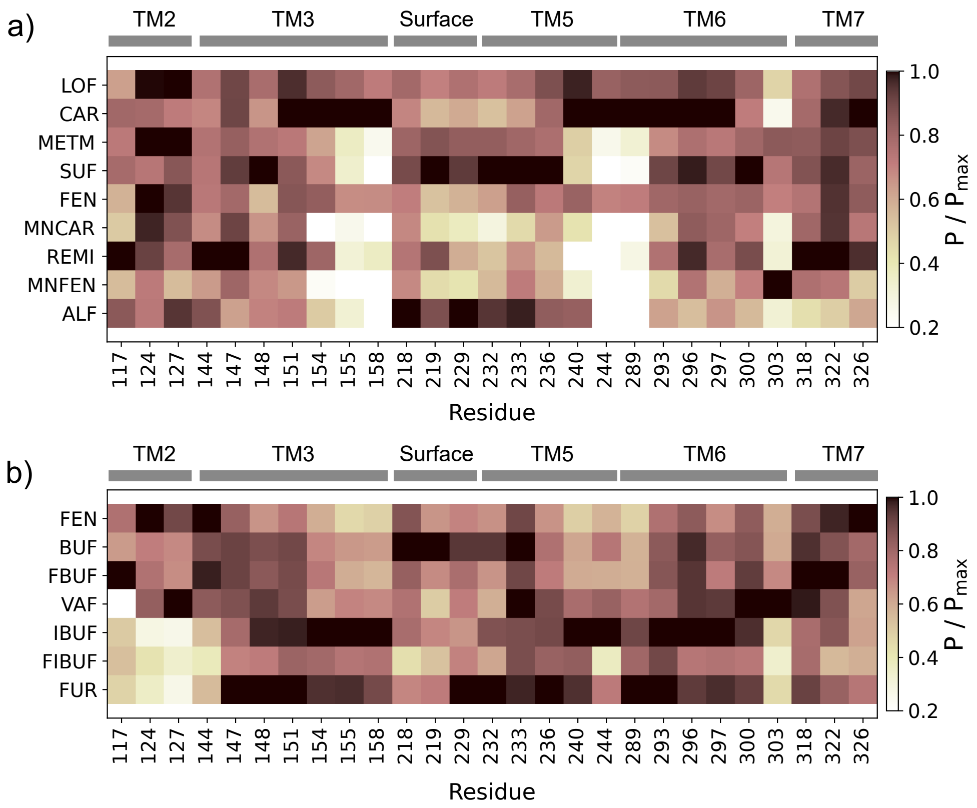


**Figure S4: Interactions between fentanyls and the mOR residues. a)** Contact profiles for fentanyls with R4 and/or R1 modification with the relevant mOR residues. **b)** Contact profiles for fentanyls with R2 modification with the relevant mOR residues. $P$ is the fraction of the analyzed trajectories that a compound interacted with a residue defined by a heavy-atom distance cutoff of 5 Å. $P_{\mathrm{max}}$ is the maximum value of contact fractions ($P$) among all compounds.

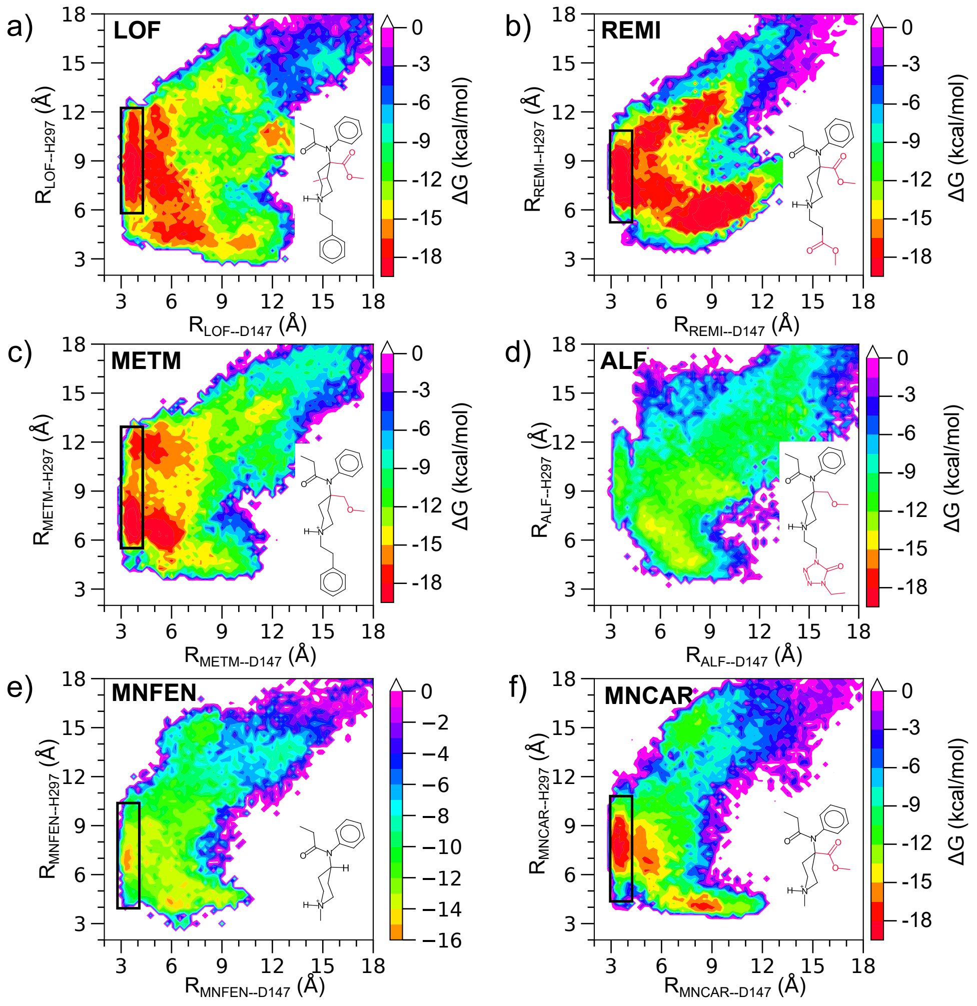


**Figure S5: Approximate free energy surfaces illustrating the D147 salt bridge and the H297 hydrogen bond with the rest of fentanyls with R4 and/or R1 modification. a–f)** $R_{piper\text{-}D147}$, which is the distance between the piperidine nitrogen and carboxylate carbon atom of D147, illustrates the formation of the D147 salt bridge. $R_{piper\text{-}H297}$, which is the distance between the piperidine nitrogen and the nearest imidazole nitrogen of H297, illustrates the formation of the H297 hydrogen bond. The approximate free energies were calculated using the reweighting protocol[S35] implemented in PLUMED (version 2.5).[S23]

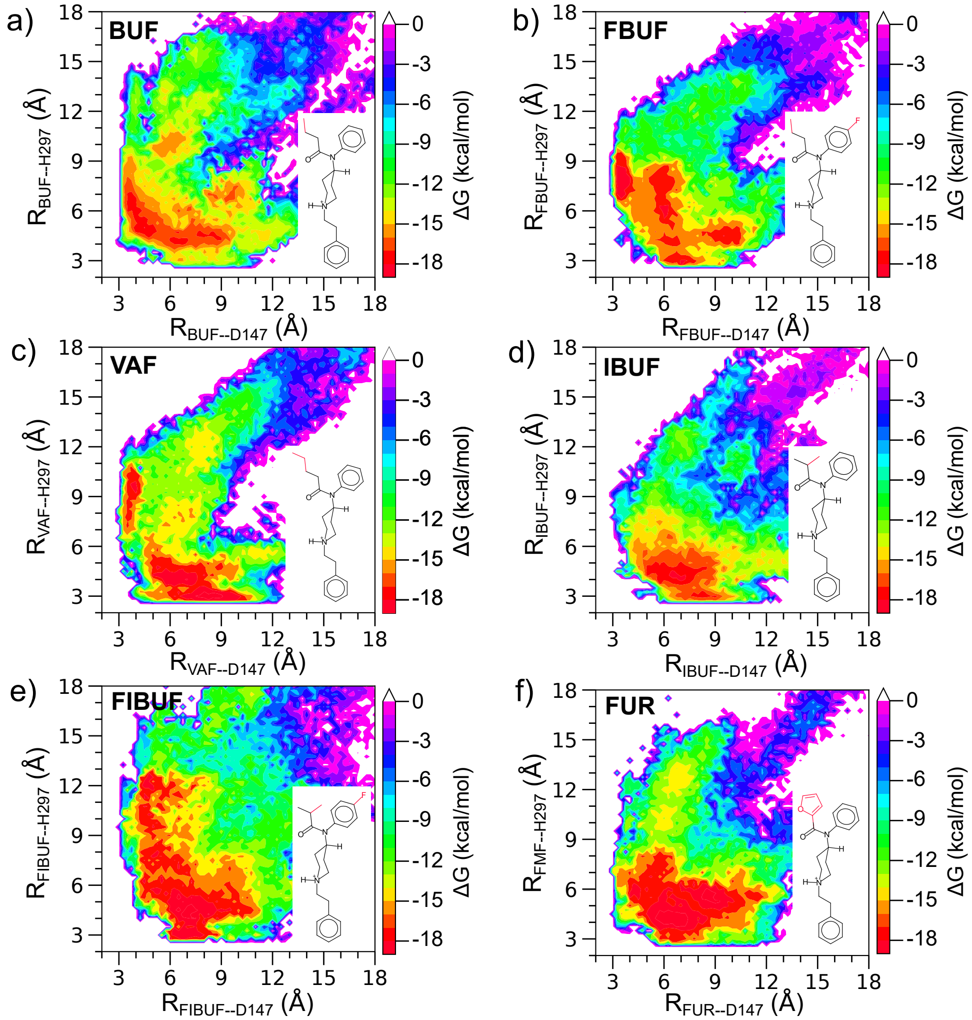


**Figure S6: Approximate free energy surfaces illustrating the D147 salt bridge and the H297 hydrogen bond with the rest of fentanyls with R2 modification. a–f)** The free energy surfaces are projected onto two distances, $R_{piper\text{-}D147}$ and $R_{piper\text{-}H297}$. The distances are defined in SFig. S5. The approximate free energies were calculated using the reweighting protocol[S35] implemented in PLUMED (version 2.5).[S23]

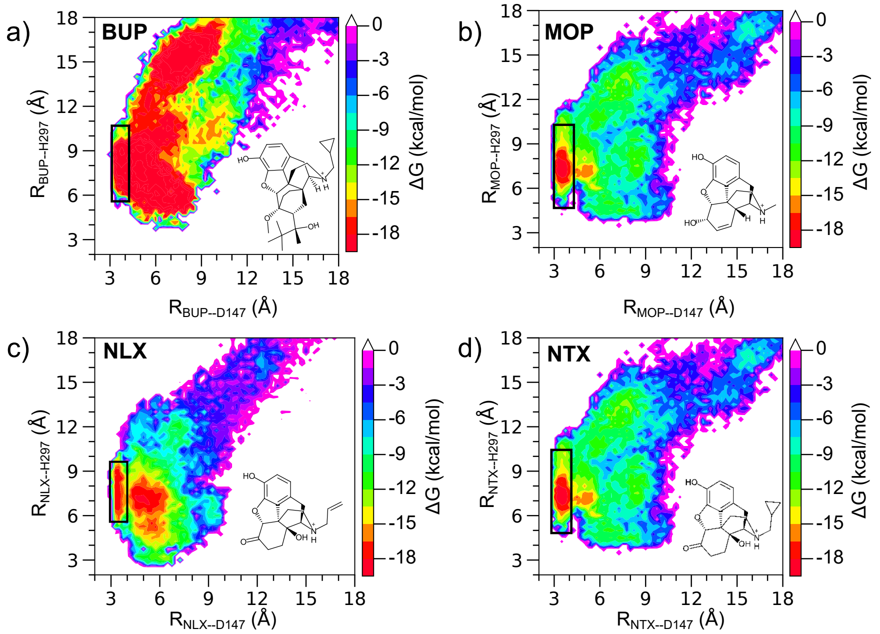


**Figure S7: The amine-D147 salt bridge and amine-H297 h-bond interaction for morphinans.** Approximate free energy surface of BUP (a), MOP (b), NLX (c) and NTX (d) projected onto two distances, $R_{amine\text{–}D147}$ and $R_{amine\text{–}H297}$. The distances are defined in Fig. S5. The approximate free energies were calculated using the reweighting protocol[S35] implemented in PLUMED (version 2.5).[S23]

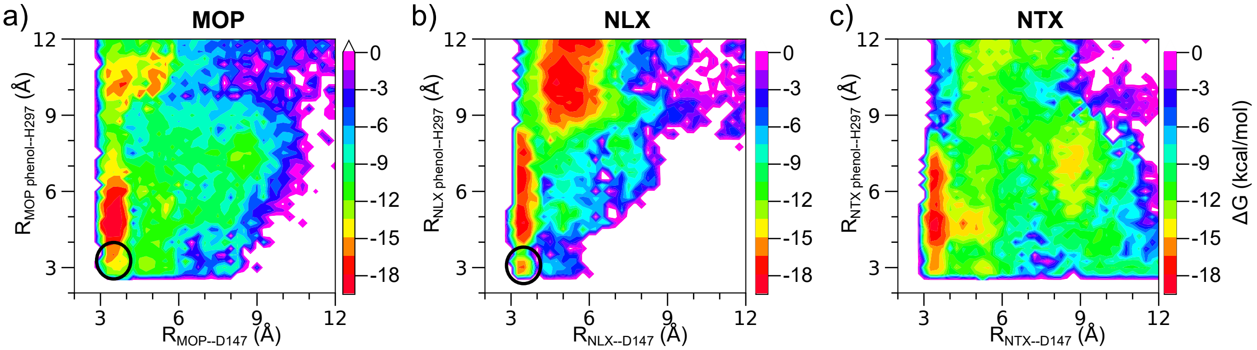


**Figure S8: The piperidine amine-D147 salt bridge and phenol-H297 h-bond interactions of the morphinans.** The approximate free energy surface of MOP (a), NLX (b) and NTX (b) projected onto $R_{\text{amine-D147}}$ and $R_{\text{phenol-H297}}$. $R_{\text{amine-D147}}$ is defined in Fig. S5. $R_{\text{Phenol-H297}}$ is defined as the distance between the phenolic oxygen and the nearest imidazole nitrogen of H297. Circles indicate the formation of both amine–D147 salt bridge and phenol–H297 h-bond. The approximate free energies were calculated using the reweighting protocol[S35] implemented in PLUMED (version 2.5).[S23,S35]

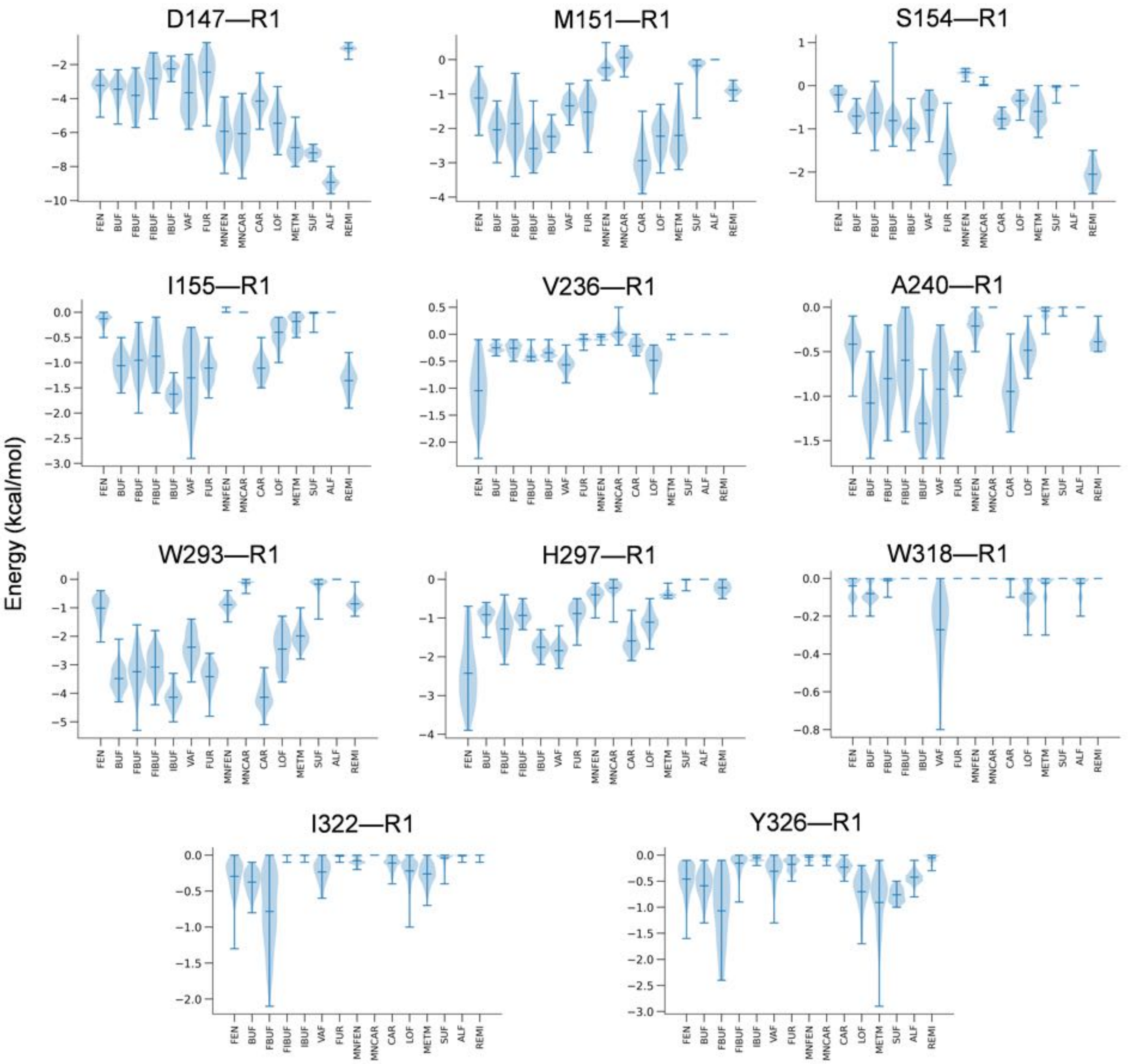


**Figure S9:** Violin plots of the reweighted interaction energies between selected mOR residues and the R1 group of different fentanyls. The maximum, median, and minimum are indicated by upper, middle and lower bars, respectively.

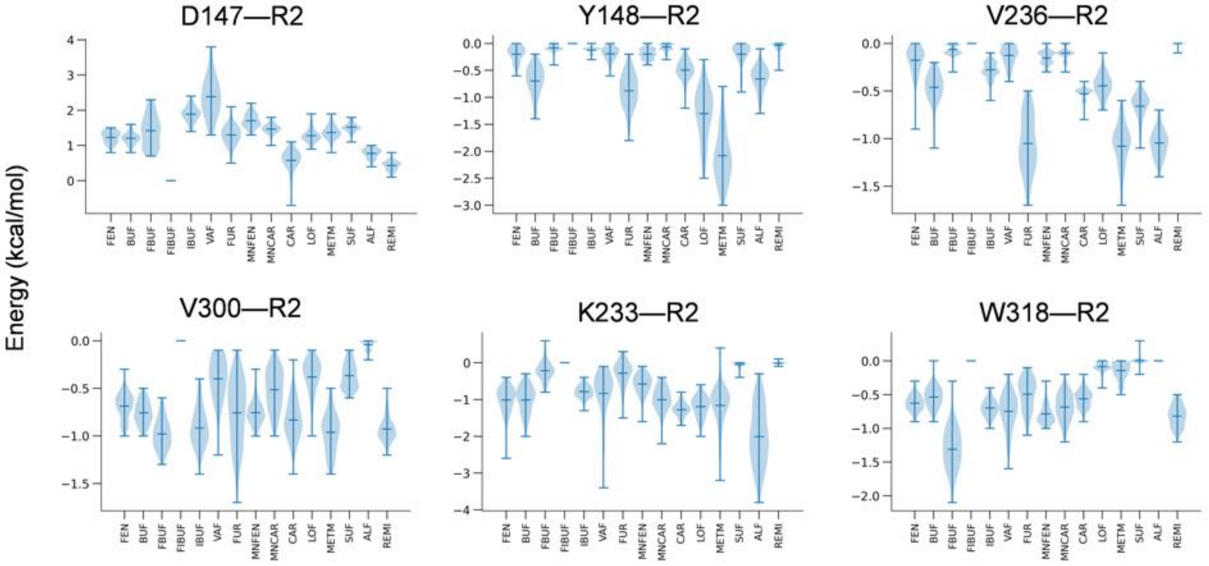


**Figure S10:** Violin plots of the reweighted interaction energies between the selected mOR residues and the R2 group of different fentanyls. The maximum, median, and minimum are indicated by upper, middle and lower bars, respectively.

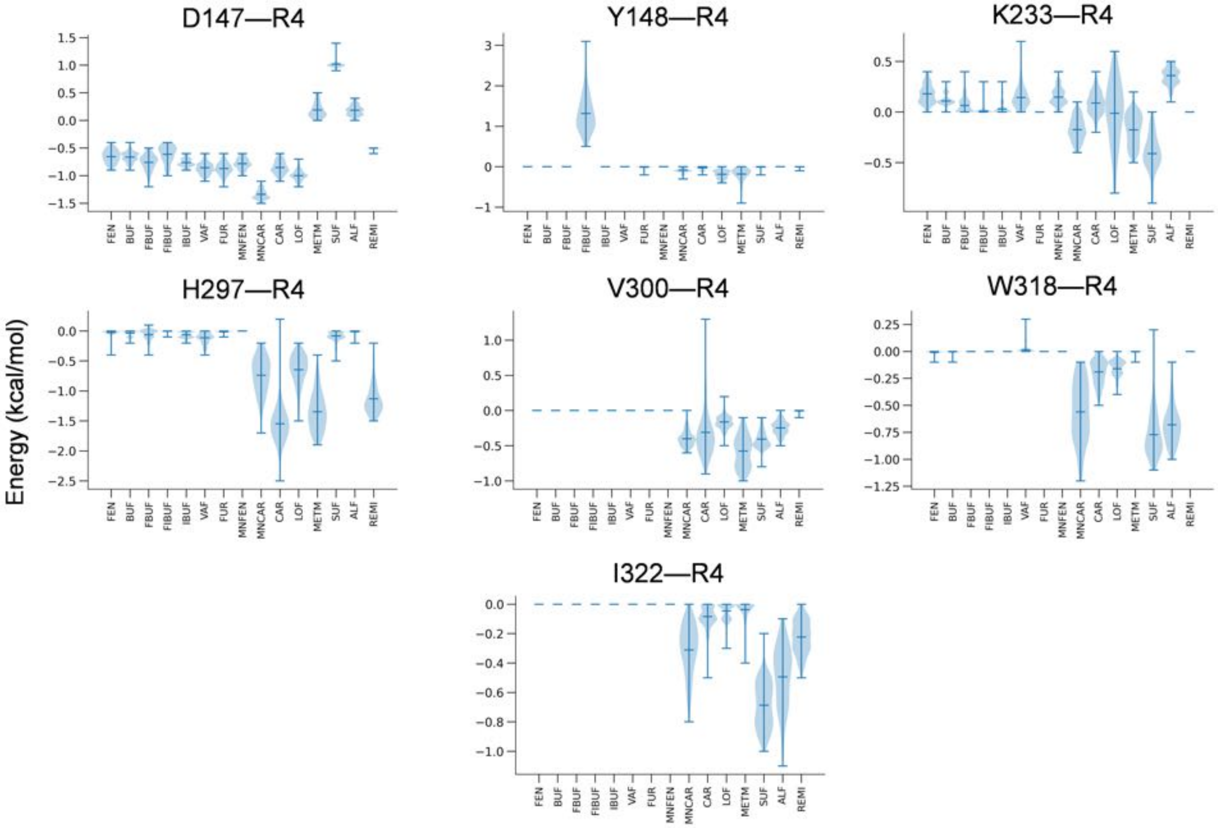


**Figure S11:** Violin plots of the reweighted interaction energies between the selected mOR residues and the R4 group of different fentanyls. The maximum, median, and minimum are indicated by upper, middle and lower bars, respectively.

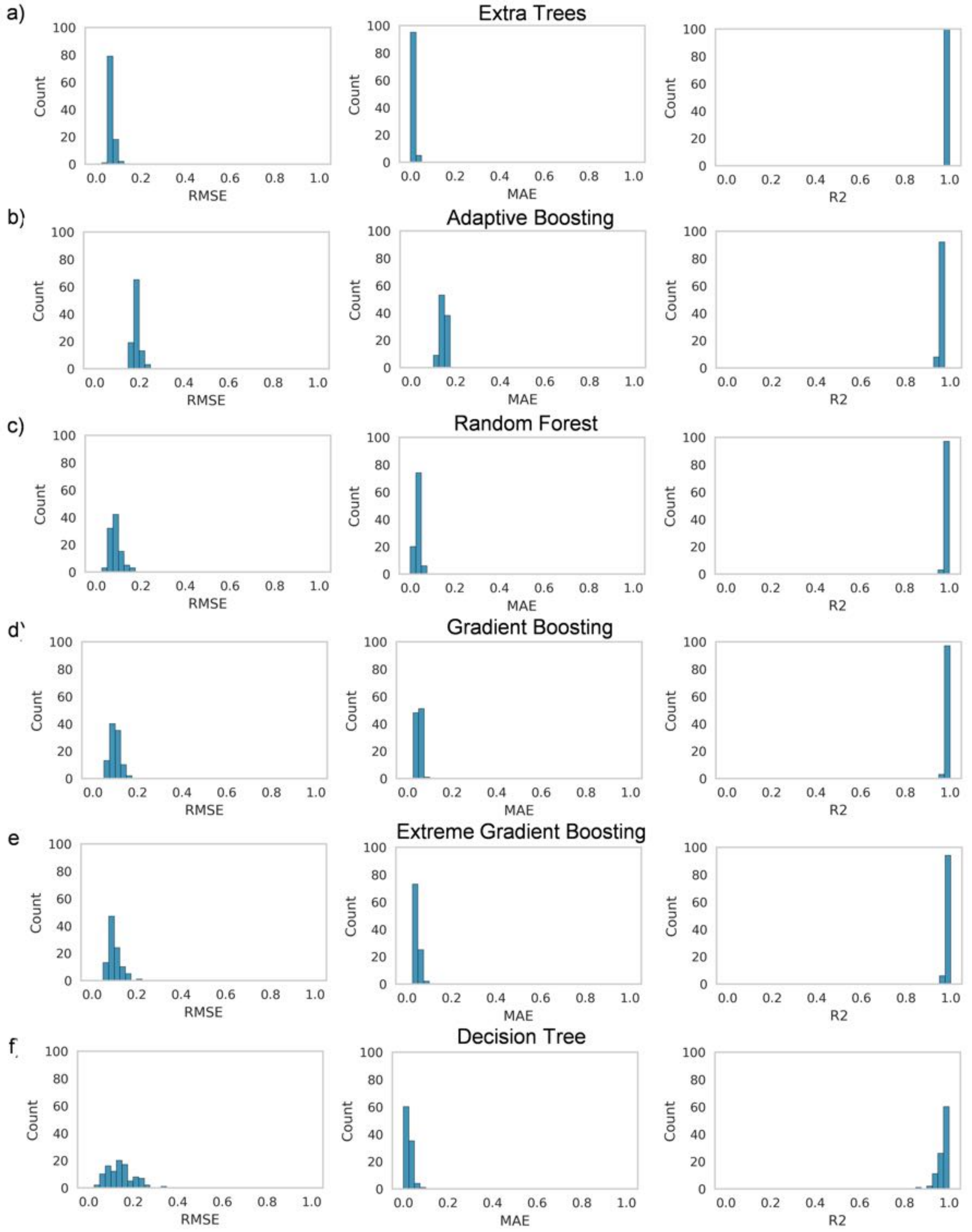


**Figure S12: Model qualities of the tree-based regression models (all 24 features).** Histograms of the root-mean squared error (RMSE), mean absolute error (MAE), and $R^2$ values were calculated over 100 independent ML trials.

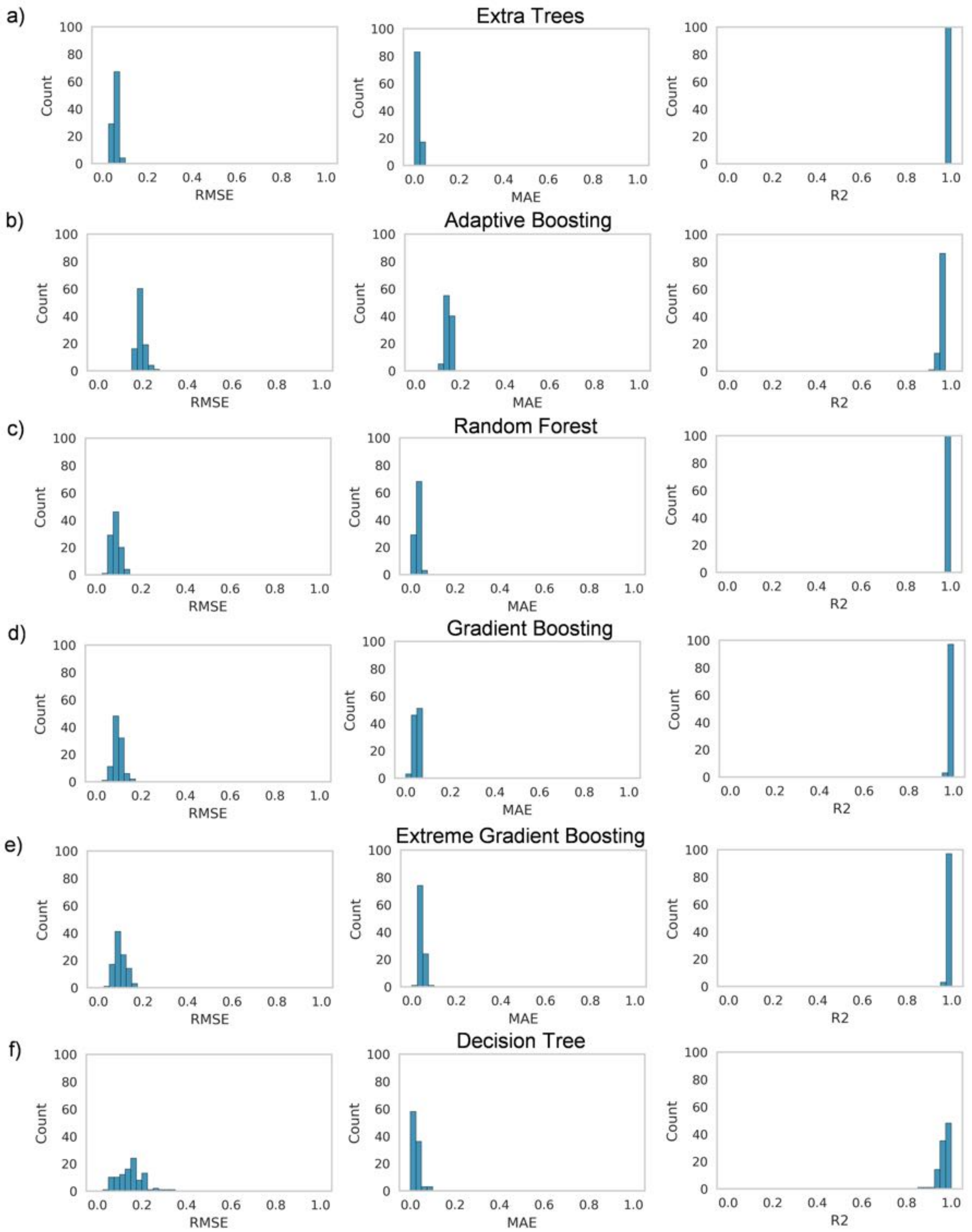


**Figure S13: Model qualities of the tree-based regression models (15 features).** Histograms of the root-mean squared error (RMSE), mean absolute error (MAE), and $R^2$ values were calculated over 100 independent ML trials. The 15 features are the D147-R1, M151-R1, S154-R1, V236-R1, H297-R1, W293-R1, Y326-R1, D147-R2, Y148-R2, V236-R2, D147-R4, K233-R4, H297-R4, V300-R4 pairs.

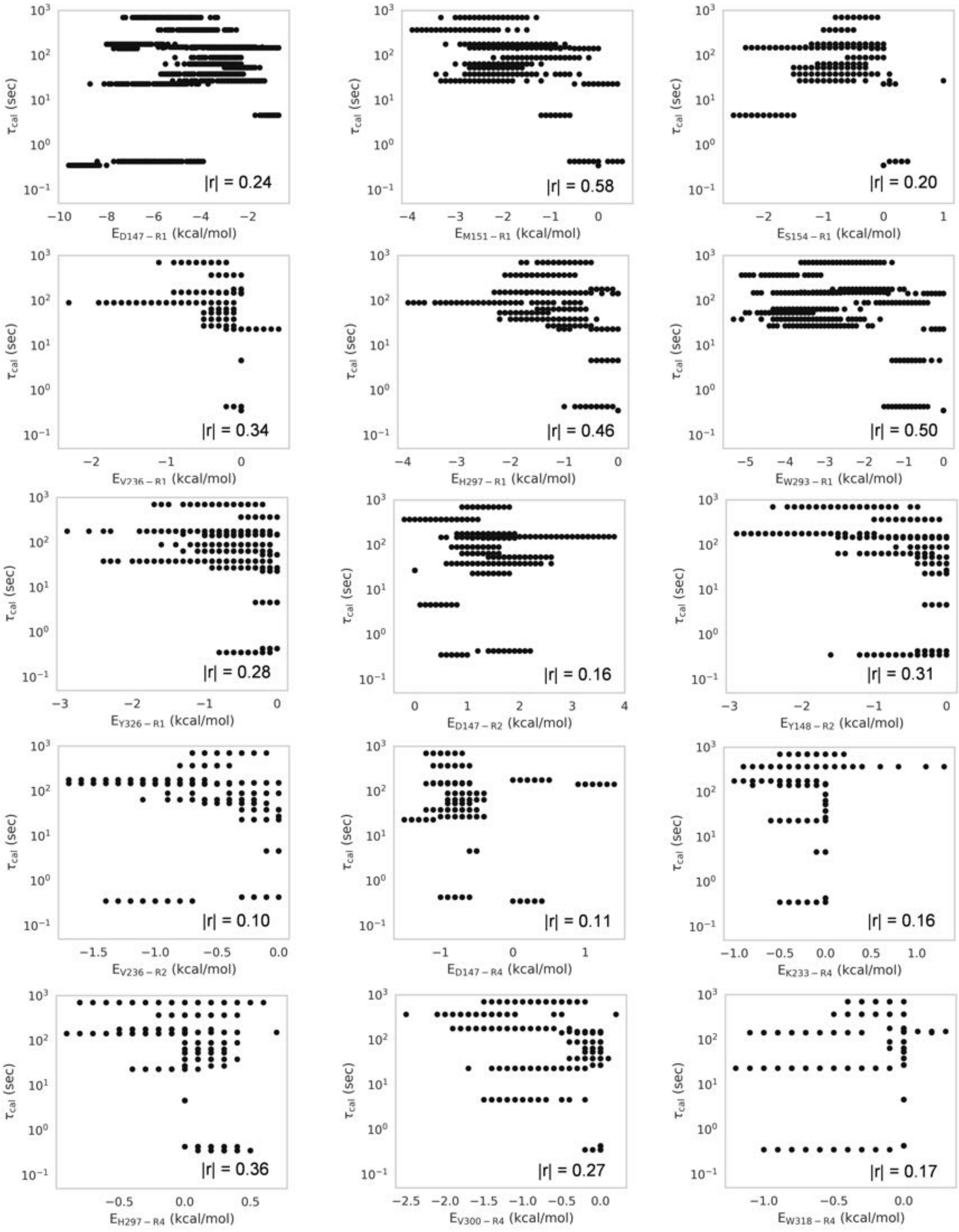


**Figure S14: Correlation between the calculated residence times ($\tau_{\mathrm{cal}}$) and the residue-substituent interaction energies.** The black dots correspond to the average interaction energies for the individual compounds. For each compound, the energies were randomly sampled 100 times, each of which comprises 2,000 snapshots randomly selected from the frames of the combined trajectories. Note, some of the energy values overlap.

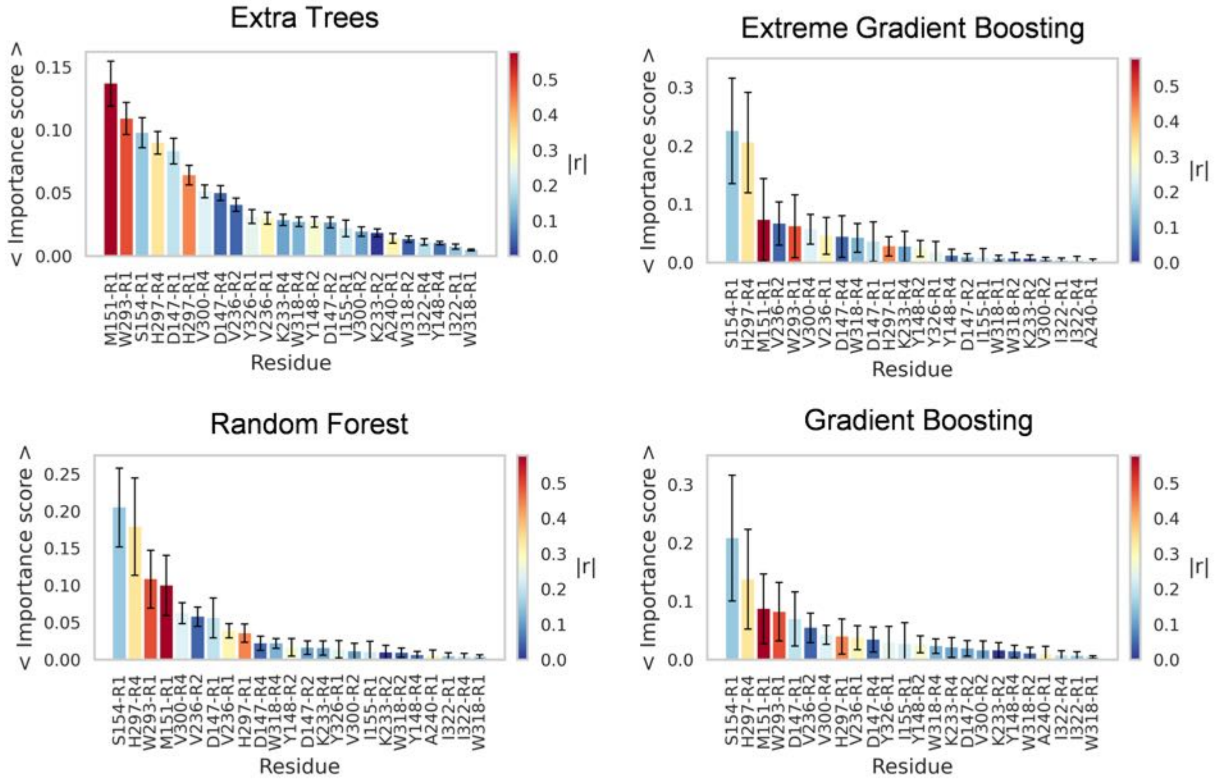


**Figure S15: Average feature importance scores for the top four models (all 24 features).** The feature importance scores shown are the averages of the mean decrease impurity importance scores calculated over 100 trials. The error bars correspond to the standard deviations. The feature are color coded based on the Pearson's correlation coefficient with the residence times.

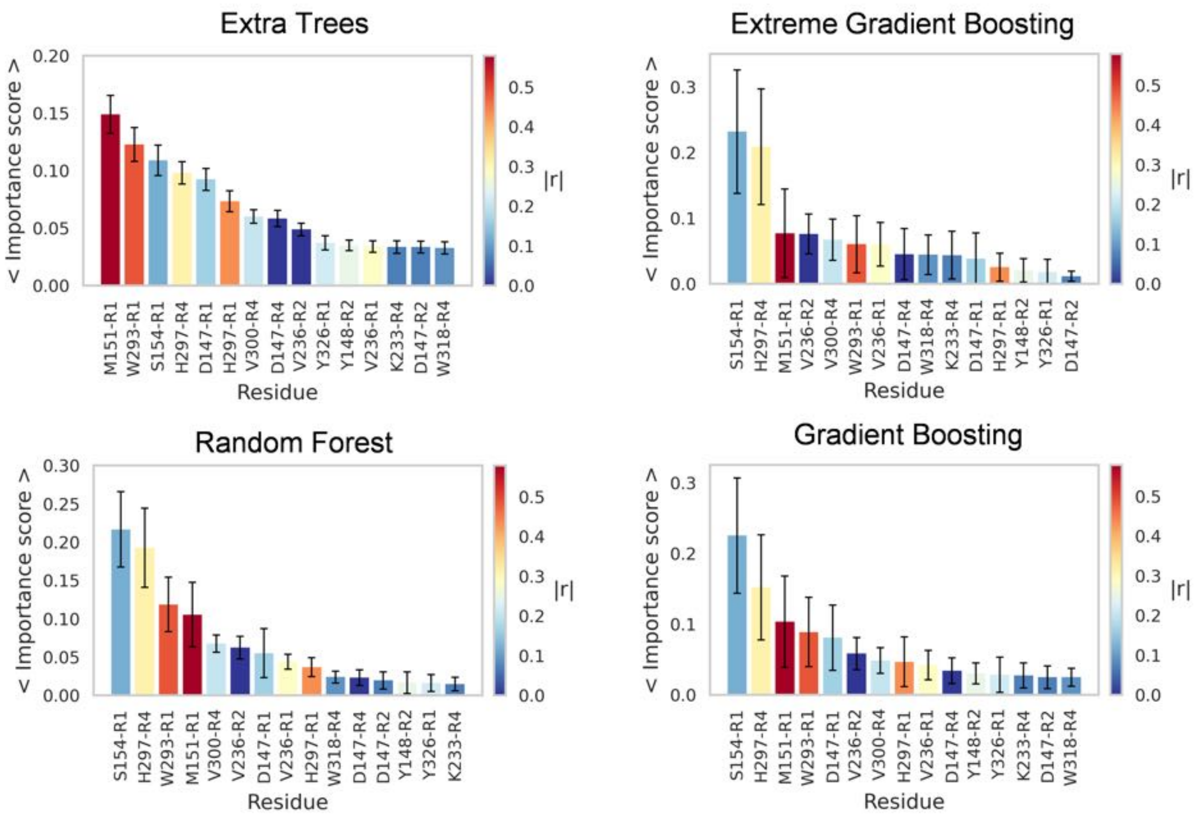


**Figure S16: Average feature importance scores for the top four models (top 15 features).** The feature importance scores shown are the averages of the mean decrease impurity importance scores calculated over 100 trials. The error bars correspond to the standard deviations. The feature are color coded based on the Pearson's correlation coefficient with the residence times.

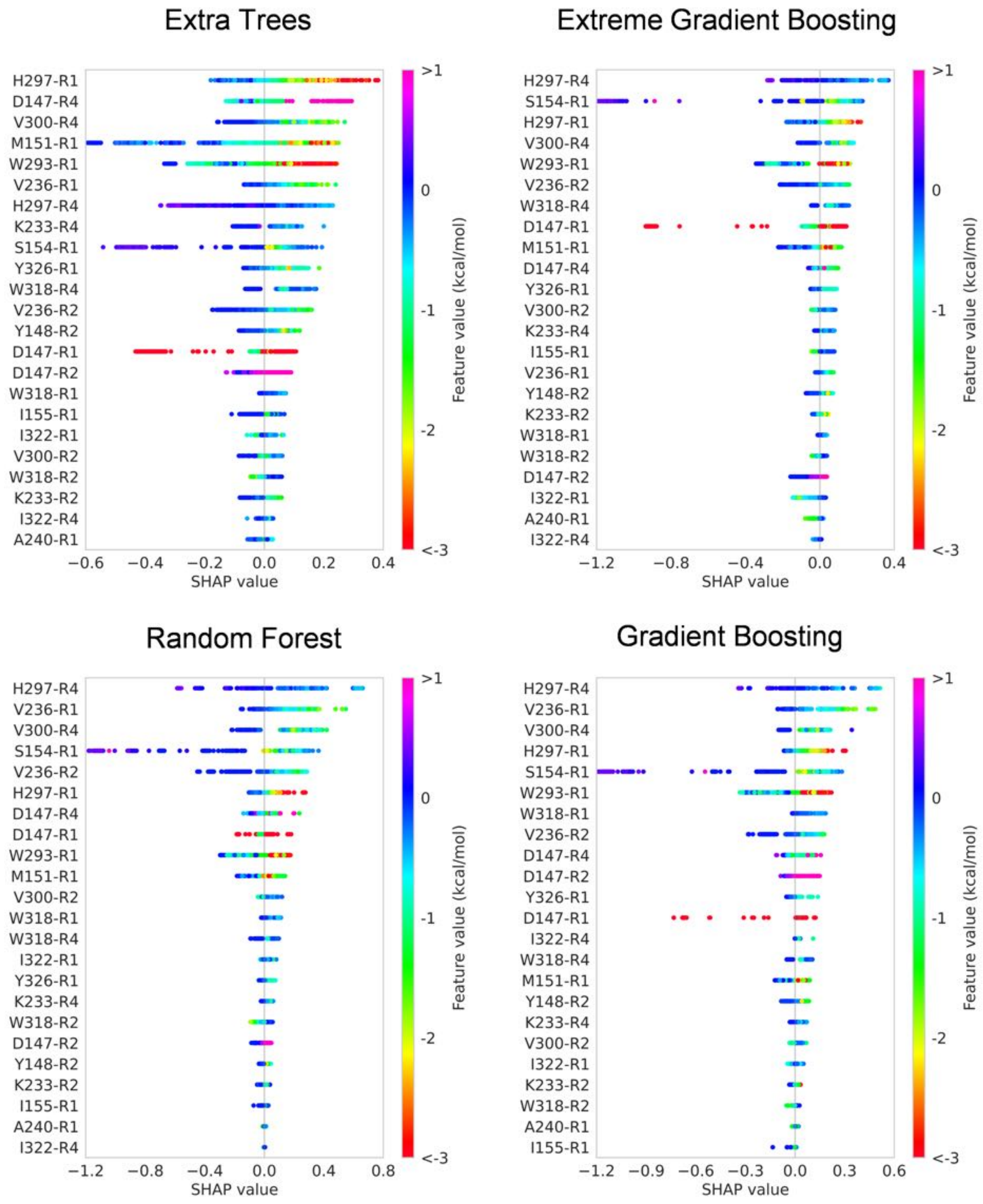


**Figure S17: Beeswarm plots of the SHAP values for the features used in the top four models (all 24 features).** The bee swarm plots combined all 1,500 data points. The SHAP values represent the impact of the individual interactions on the residence times. Color indicates the feature value (energy in kcal/mol).

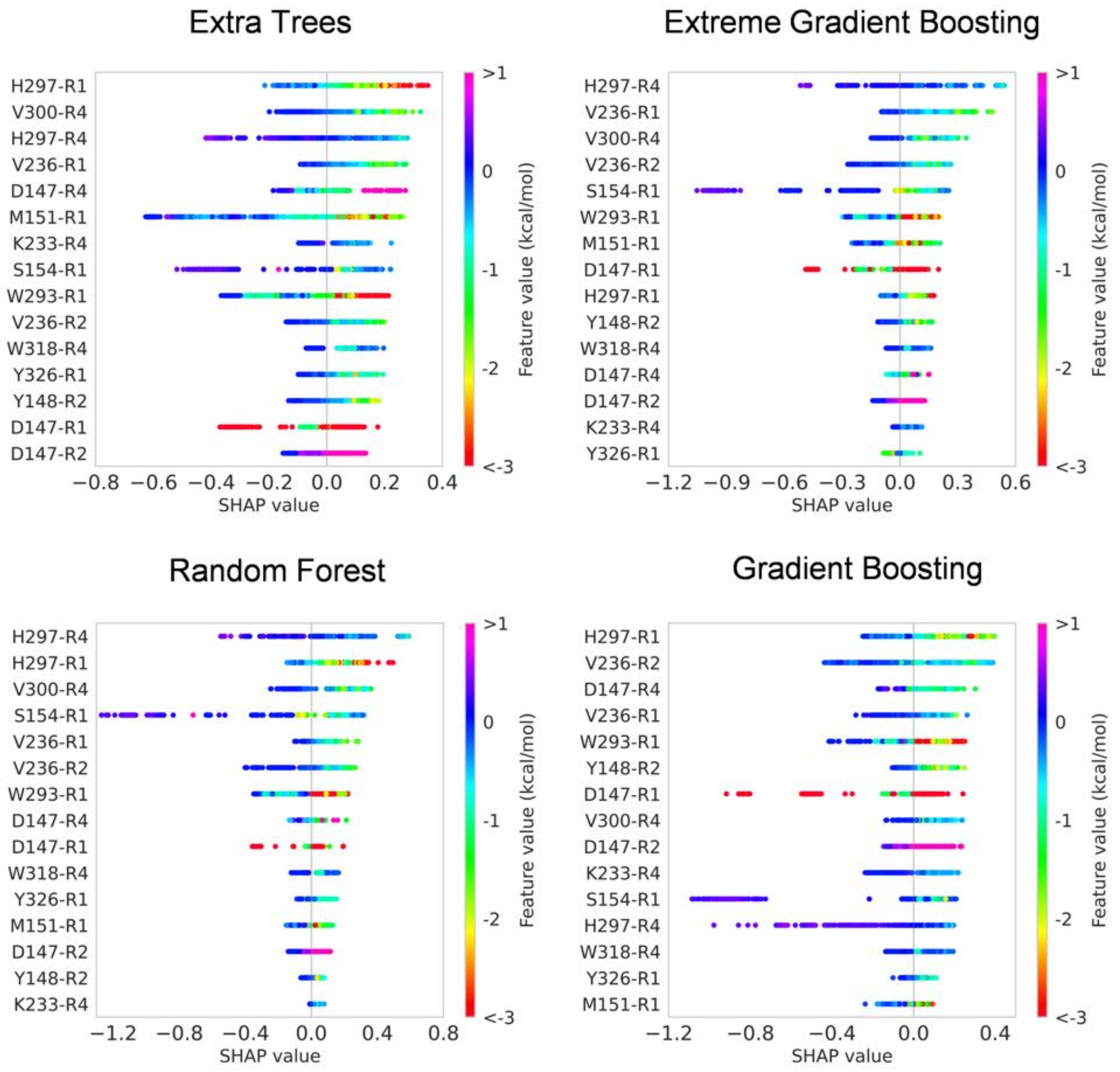


**Figure S18: Beeswarm plots of the SHAP values for the features used in the top four models. (top 15 features)** The bee swarm plots combined all 1,500 data points. The SHAP values represent the impact of the individual interactions on the residence times. Color indicates the feature value (energy in kcal/mol). The features were ranked (from top to bottom) based on the maximum impact on the prediction outcome (maximum value of the SHAP values).

### References


(S1) Bajusz, D.; Rácz, A.; Héberger, K. Why Is Tanimoto Index an Appropriate Choice for Fingerprint-Based Similarity Calculations? *J Cheminform* **2015**, *7*, 20.

(S2) Landrum, G. A. RDKit: Open-source cheminformatics.

(S3) Rogers, D.; Hahn, M. Extended-Connectivity Fingerprints. *J. Chem. Inf. Model.* **2010**, *50*, 742–754.

(S4) Maggiora, G.; Vogt, M.; Stumpfe, D.; Bajorath, J. Molecular similarity in medicinal chemistry. *J. Med. Chem.* **2014**, *57*, 3186–3204.

(S5) Mahinthichaichan, P.; Vo, Q. N.; Ellis, C. R.; Shen, J. Kinetics and Mechanism of Fentanyl Dissociation from the $\mu$-Opioid Receptor. *JACS Au* **2021**, *1*, 2208–2215.

(S6) Vo, Q. N.; Mahinthichaichan, P.; Shen, J.; Ellis, C. R. How $\mu$-Opioid Receptor Recognizes Fentanyl. *Nat Commun* **2021**, *12*, 984.

(S7) Huang, W.; Manglik, A.; Venkatakrishnan, A. J.; Laeremans, T.; Feinberg, E. N.; Sanborn, A. L.; Kato, H. E.; Livingston, K. E.; Thorsen, T. S.; Kling, R. C.; Granier, S.; Gmeiner, P.; Husbands, S. M.; Traynor, J. R.; Weis, W. I.; Steyaert, J.; Dror, R. O.; Kobilka, B. K. Structural Insights into $\mu$-Opioid Receptor Activation. *Nature* **2015**, *524*, 315–321.

(S8) Manglik, A.; Kruse, A. C.; Kobilka, T. S.; Thian, F. S.; Mathiesen, J. M.; Sunahara, R. K.; Pardo, L.; Weis, W. I.; Kobilka, B. K.; Granier, S. Crystal Structure of the $M$-Opioid Receptor Bound to a Morphinan Antagonist. *Nature* **2012**, *485*, 321–326.

(S9) Wallace, J. A.; Shen, J. K. Continuous Constant pH Molecular Dynamics in Explicit Solvent with pH-Based Replica Exchange. *J. Chem. Theory Comput.* **2011**, *7*, 2617–2629.

(S10) Huang, Y.; Chen, W.; Dotson, D. L.; Beckstein, O.; Shen, J. Mechanism of pH-Dependent Activation of the Sodium-Proton Antiporter NhaA. *Nat. Commun.* **2016**, *7*, 12940.

(S11) Huang, Y.; Henderson, J. A.; Shen, J. Continuous constant pH Molecular Dynamics Simulations of Transmembrane Proteins. *Methods Mol. Biol.* **2021**, in press.

(S12) Phillips, J. C.; Braun, R.; Wang, W.; Gumbart, J.; Tajkhorshid, E.; Villa, E.; Chipot, C.; Skeel, R. D.; Kalé, L.; Schulten, K. Scalable Molecular Dynamics with NAMD. *J. Comput. Chem.* **2005**, *26*, 1781–1802.

(S13) Best, R. B.; Zhu, X.; Shim, J.; Lopes, P. E. M.; Mittal, J.; Feig, M.; MacKerell, A. D. Optimization of the Additive CHARMM All-Atom Protein Force Field Targeting Improved Sampling of the Backbone $\phi$, $\psi$ and Side-Chain $\chi_1$ and $\chi_2$ Dihedral Angles. *J. Chem. Theory Comput.* **2012**, *8*, 3257–3273.

(S14) Klauda, J. B.; Venable, R. M.; Freites, J. A.; O'Connor, J. W.; Tobias, D. J.; Mondragon-Ramirez, C.; Vorobyov, I.; MacKerell, A. D.; Pastor, R. W. Update of the CHARMM All-Atom Additive Force Field for Lipids: Validation on Six Lipid Types. *J. Phys. Chem. B* **2010**, *114*, 7830–7843.

(S15) MacKerell, A. D.; Bashford, D.; Bellott, M.; Dunbrack, R. L.; Evanseck, J. D.; Field, M. J.; Fischer, S.; Gao, J.; Guo, H.; Ha, S.; Joseph-McCarthy, D.; Kuchnir, L.; Kuczera, K.; Lau, F. T. K.; Mattos, C.; Michnick, S.; Ngo, T.; Nguyen, D. T.; Prodhom, B.; Reiher, W. E.; Roux, B.; Schlenkrich, M.; Smith, J. C.; Stote, R.; Straub, J.; Watanabe, M.; Wiórkiewicz-Kuczera, J.; Yin, D.; Karplus, M. All-Atom Empirical Potential for Molecular Modeling and Dynamics Studies of Proteins †. *J Phys Chem B* **1998**, *102*, 3586–3616.

(S16) Vanommeslaeghe, K.; MacKerell, A. D. Automation of the CHARMM General Force

Field (CGenFF) I: Bond Perception and Atom Typing. *J Chem Inf Model* **2012**, *52*, 3144–3154.

(S17) Vanommeslaeghe, K.; Raman, E. P.; MacKerell, A. D. Automation of the CHARMM General Force Field (CGenFF) II: Assignment of Bonded Parameters and Partial Atomic Charges. *J Chem Inf Model* **2012**, *52*, 3155–3168.

(S18) Ryckaert, J.-P.; Ciccotti, G.; Berendsen, H. J. Numerical Integration of the Cartesian Equations of Motion of a System with Constraints: Molecular Dynamics of n-Alkanes. *J. Comput. Phys.* **1977**, *23*, 327–341.

(S19) Martyna, G. J.; Tobias, D. J.; Klein, M. L. Constant Pressure Molecular Dynamics Algorithms. *J. Chem. Phys.* **1994**, *101*, 4177–4189.

(S20) Feller, S. E.; Zhang, Y.; Pastor, R. W.; Brooks, B. R. Constant Pressure Molecular Dynamics Simulation: The Langevin Piston Method. *J. Chem. Phys.* **1995**, *103*, 4613–4621.

(S21) Darden, T.; York, D.; Pedersen, L. Particle Mesh Ewald: An $N \cdot \log(N)$ Method for Ewald Sums in Large Systems. *J. Chem. Phys.* **1993**, *98*, 10089–10092.

(S22) Humphrey, W.; Dalke, A.; Schulten, K. VMD: Visual Molecular Dynamics. *J. Mol. Graph.* **1996**, *14*, 33–38.

(S23) Tribello, G. A.; Bonomi, M.; Branduardi, D.; Camilloni, C.; Bussi, G. PLUMED 2: New Feathers for an Old Bird. *Comput. Phys. Commun.* **2014**, *185*, 604–613.

(S24) Fiorin, G.; Klein, M. L.; Hénin, J. Using Collective Variables to Drive Molecular Dynamics Simulations. *Mol. Phys.* **2013**, *111*, 3345–3362.

(S25) Laio, A.; Parrinello, M. Escaping Free-Energy Minima. *Proc. Natl. Acad. Sci.* **2002**, *99*, 12562–12566.

(S26) Barducci, A.; Bussi, G.; Parrinello, M. Well-Tempered Metadynamics: A Smoothly Converging and Tunable Free-Energy Method. *Phys. Rev. Lett.* **2008**, *100*, 020603.

(S27) Casasnovas, R.; Limongelli, V.; Tiwary, P.; Carloni, P.; Parrinello, M. Unbinding Kinetics of a P38 MAP Kinase Type II Inhibitor from Metadynamics Simulations. *J. Am. Chem. Soc.* **2017**, *139*, 4780–4788.

(S28) Lamim Ribeiro, J. M.; Provasi, D.; Filizola, M. A Combination of Machine Learning and Infrequent Metadynamics to Efficiently Predict Kinetic Rates, Transition States, and Molecular Determinants of Drug Dissociation from G Protein-Coupled Receptors. *J. Chem. Phys.* **2020**, *153*, 124105.

(S29) Salvalaglio, M.; Tiwary, P.; Parrinello, M. Assessing the Reliability of the Dynamics Reconstructed from Metadynamics. *J. Chem. Theory Comput.* **2014**, *10*, 1420–1425.

(S30) Palacio-Rodriguez, K.; Vroylandt, H.; Stelzl, L. S.; Pietrucci, F.; Hummer, G.; Cossio, P. Transition Rates and Efficiency of Collective Variables from Time-Dependent Biased Simulations. *J. Phys. Chem. Lett.* **2022**, *13*, 7490–7496.

(S31) Ali, M. PyCaret: An open source, low-code machine learning library in Python. 2020; PyCaret version 1.0.

(S32) Mann, J.; Samieegohar, M.; Chaturbedi, A.; Zirkle, J.; Han, X.; Ahmadi, S. F.; Eshleman, A.; Janowsky, A.; Wolfrum, K.; Swanson, T.; Bloom, S.; Dahan, A.; Olofsen, E.; Florian, J.; Strauss, D. G.; Li, Z. Development of a Translational Model to Assess the Impact of Opioid Overdose and Naloxone Dosing on Respiratory Depression and Cardiac Arrest. *Clin. Pharmacol. Therapeut.* **2022**, *112*, 1020–1032.

(S33) Pedersen, M. F.; Wróbel, T. M.; Märcher-Rørsted, E.; Pedersen, D. S.; Møller, T. C.; Gabriele, F.; Pedersen, H.; Matosiuk, D.; Foster, S. R.; Bouvier, M.; Bräuner-Osborne, H. Biased Agonism of Clinically Approved $\mu$-Opioid Receptor Agonists and TRV130 Is Not Controlled by Binding and Signaling Kinetics. *Neuropharmacol.* **2020**, *166*, 107718.

(S34) Cassel, J. A.; Daubert, J. D.; DeHaven, R. N. [3H]Alvimopan Binding to the $\mu$ Opioid Receptor: Comparative Binding Kinetics of Opioid Antagonists. *Eur. J. Pharmacol.* **2005**, *520*, 29–36.

(S35) Bonomi, M.; Barducci, A.; Parrinello, M. Reconstructing the Equilibrium Boltzmann Distribution from Well-Tempered Metadynamics. *J. Comput. Chem.* **2009**, *30*, 1615–1621.
